# Subsynaptic mobility of presynaptic mGluR types is differentially regulated by intra- and extracellular interactions

**DOI:** 10.1101/2020.07.06.188995

**Authors:** Anna Bodzęta, Florian Berger, Harold D. MacGillavry

## Abstract

Presynaptic metabotropic glutamate receptors (mGluRs) are essential for activity-dependent modulation of synaptic transmission. However, the mechanisms that control the subsynaptic dynamics of these receptors and their contribution to synaptic signaling are poorly understood. Here, using complementary super-resolution microscopy and single-molecule tracking techniques, we provide novel insights into the molecular mechanisms that control the nanoscale distribution and mobility of presynaptic mGluRs in hippocampal neurons. We demonstrate that the group II receptor mGluR2 localizes diffusely along the axon and boutons, and is highly mobile, while the group III receptor mGluR7 is stably anchored at the active zone, indicating that distinct mechanisms underlie the dynamic distribution of these receptor types. Surprisingly, using domain swapping experiments we found that intracellular interactions modulate surface diffusion of mGluR2, but not mGluR7. Instead, we found that immobilization of mGluR7 at the active zone relies on its extracellular domain. Importantly, receptor activation or increase in synaptic activity did not alter the surface mobility of presynaptic mGluRs. Finally, computational modeling of presynaptic mGluR activity revealed that the precise subsynaptic distribution of mGluRs controls their activation probability and thus directly impacts their ability to modulate neurotransmitter release. Altogether, this study demonstrates that distinct mechanisms control surface mobility of presynaptic mGluRs to differentially contribute to the regulation of glutamatergic synaptic transmission.

## INTRODUCTION

Activity-directed modulation of synaptic efficacy underlies the ability of neuronal networks to process and store information. Presynaptic mechanisms that impinge on the neurotransmitter release machinery are a critical factor in fine-tuning synaptic efficacy. In particular, presynaptic metabotropic glutamate receptors (mGluRs) are essential negative-feedback control elements that modulate transmission by dampening glutamate release (Reiner & Levitz, 2018; Pinheiro & Mulle, 2008). Disruptions in these receptor systems severely deregulate synaptic function and specific forms of synaptic plasticity, and aberrant mGluR function has been associated with several neurological disorders such as anxiety, epilepsy, and schizophrenia, further highlighting their physiological importance (Muly *et al*, 2007; Sansig *et al*, 2001; Woolley *et al*, 2008). Nevertheless, it remains poorly understood how these receptors are organized at presynaptic sites to efficiently modulate transmission.

The eight known mGluRs (mGluR1 – mGluR8) belong to the class C G-protein-coupled receptors (GPCRs). These GPCRs exist as constitutive dimers and have distinctive large extracellular domains (ECD) that contain the ligand-binding domain connected to the prototypical 7-helix transmembrane domain (TMD) via a cysteine-rich domain. mGluRs are further divided into three groups based on their sequence homology, downstream signaling partners, and agonist selectivity (Niswender & Conn, 2010). These functionally diverse groups are expressed throughout the central nervous system but are generally targeted to specific subcellular locations. Group I mGluRs (mGluR1/5) are primarily expressed at postsynaptic sites, group II mGluRs (mGluR2/3) are present at both pre- and postsynaptic sites, and group III mGluRs (mGluR4, mGluR6-8) are located almost exclusively at presynaptic sites, except mGluR6 which is located at the postsynaptic site in retina bipolar cells (Petralia *et al*, 1996; Shigemoto *et al*, 1996). The presynaptic group II and III mGluRs mGluR2 and mGluR7 are both abundantly expressed in the hippocampus (Kinoshita *et al*, 1998), share substantial homology (∼60%), and both couple to inhibitory G-proteins (Gα_i/o_) that repress adenylyl cyclase activity. Nevertheless, these receptors differ significantly in their pharmacological characteristics and interactome, conferring functionally distinct roles to these receptors in synaptic transmission and plasticity.

Generally, activation of presynaptic mGluRs depresses synaptic transmission via inhibition of voltage-gated Ca^2+^-channels (VGCC), activation of K^+^ channels, or by directly modulating components of the release machinery such as Munc13, Munc18, and RIM-1 (de Jong & Verhage, 2009; Pinheiro & Mulle, 2008). As such, these receptors have been implicated in the regulation of both short-term plasticity as well as long-term depression of synaptic responses (Martín *et al*, 2007; Millán *et al*, 2002; Robbe *et al*, 2002; Kamiya & Ozawa, 1999; Okamoto *et al*, 1994; Pelkey *et al*, 2008, 2005). However, signaling events downstream of presynaptic mGluRs can also potentiate release, and particularly mGluR7 has been postulated to bidirectionally regulate synaptic transmission (Martín *et al*, 2018, 2010; Dasgupta *et al*, 2020; Klar *et al*, 2015). Thus, presynaptic mGluRs modulate synaptic transmission through a variety of downstream effectors, and the functional outcome of mGluR activation is probably determined by the frequency and duration of synaptic signals. Additionally, the subsynaptic distribution and dynamics of presynaptic mGluRs are likely to influence their ability to become activated and engage local downstream signaling partners. In particular, since these receptors have different affinities for glutamate, their subsynaptic position relative to the point of glutamate release ultimately determines their probability of activation. mGluR2 has a moderate to high affinity for glutamate (in the micromolar range) and its positioning relative to the release site might thus only modestly affect its contribution to regulating release probability. In contrast, when measured in non-neuronal cells, the affinity of mGluR7 for glutamate is exceptionally low, in the millimolar range (0.5-2.5 mM) (Schoepp *et al*, 1999). In addition, mGluRs are obligatory dimers and activation of single subunits in an mGluR dimer produces only low-efficacy activation. Given that release events produce only brief, 1-3 mM peaks in glutamate concentration in the synaptic cleft (Lisman *et al*, 2007; Diamond & Jahr, 1997), it has thus been questioned whether mGluR7 at neuronal synapses, even when placed immediately adjacent to release sites, will ever be exposed to sufficient levels of glutamate to become activated (Pinheiro & Mulle, 2008). However, this is in contrast with the wealth of physiological evidence from different model systems that show that mGluR7 is a key modulator of synaptic transmission (Sansig *et al*, 2001; Millán *et al*, 2002; Klar *et al*, 2015; Bushell *et al*, 2002; Martín *et al*, 2018; Pelkey *et al*, 2008, 2005). Thus, the precise localization of presynaptic mGluRs determines their activation probability and greatly impacts their ability to modulate synaptic transmission through local downstream effectors. Nevertheless, quantitative insight in the dynamic distribution of presynaptic mGluRs in live neurons and the mechanisms that control their dynamic positioning is lacking.

Here, to understand how mGluR2 and mGluR7 contribute to synaptic transmission in hippocampal neurons, we studied how the dynamic positioning of the subsynaptic distribution of these receptors is mechanistically controlled. Using complementary super-resolution imaging approaches, we found that mGluR2 is highly dynamic and localized throughout the axon, while mGluR7 is immobilized at presynaptic active zones. We found that the mobility of mGluR2 is mainly mediated by its intercellular domain. Surprisingly, we found that the specific positioning of mGluR7 at the active zone is not controlled by intracellular interactions but relies on extracellular interactions. Furthermore, a computational model of mGluR activation at presynaptic sites indicates that mGluR2 activation is only loosely coupled to release site location, while activation of mGluR7 is inefficient, even when localized within a few nanometers of the release site or during high-frequency stimulation patterns. Based on our findings, we propose that the different mechanisms that control presynaptic mGluR positioning ensure the differential contribution of these receptors to transmission.

## RESULTS

### Distinct differences in the subsynaptic distribution of presynaptic mGluR subtypes

The precise spatial distribution of mGluR subtypes at presynaptic sites likely determines their functional contribution to the modulation of synaptic transmission. To compare the subsynaptic distribution of presynaptic group II and III mGluRs in hippocampal neurons, we determined the localization of mGluR2 (group II) and mGluR7 (group III) relative to the active zone marker Bassoon (Bsn) using two-color gated stimulated emission depletion (gSTED) super-resolution microscopy. To visualize mGluR2, we tagged endogenous mGluR2 with super-ecliptic pHluorin (SEP), a pH-sensitive variant of GFP, using a recently developed CRISPR/Cas9-mediated knock-in approach (Willems *et al*, 2020). Because the level of endogenous mGluR2 expression was low, we enhanced the SEP signal using anti-GFP staining to reliably measure mGluR2 distribution. We found that mGluR2 was localized both in axons and dendrites (Fig. S1A), as reported previously (Ohishi *et al*, 1994), but even though an earlier study suggested that mGluR2 is located in the preterminal region of the axon, and not in presynaptic boutons (Shigemoto *et al*, 1997), we detected mGluR2 both in the axon shaft and within synaptic boutons (Fig. 1A). However, as is apparent from line profiles of the fluorescence intensity of mGluR2 signal along Bsn-labeled puncta, the mGluR2 signal was largely excluded from presynaptic active zones (Fig. 1B). Confirming this finding, a similar distribution pattern was observed using antibody labeling for mGluR2/3 (Fig. S1B, C), further indicating that presynaptic group II mGluRs are distributed throughout the axon but excluded from active zones. Immunostaining for the group III mGluR, mGluR7 labeled a subset of neurons in our cultures (Fig. 1C, S1D), consistent with previous studies (Shigemoto *et al*, 1996; Tomioka *et al*, 2014). In contrast to mGluR2, line profiles indicated that the maximum intensity of mGluR7 labeling coincided with the Bsn-marked active zone (Fig. 1D). Co-localization analysis further confirmed this, showing that the majority of mGluR7-positive puncta overlap with Bsn-positive puncta, while mGluR2 labeling showed a striking lack of overlap with Bsn (co-localization with Bsn-positive puncta, mGluR2: 0.12 ± 0.02, mGluR7: 0.54 ± 0.03; Fig. 1E). Together, these results indicate that two presynaptic mGluR subtypes that are both implicated in the regulation of presynaptic release properties, have distinct subsynaptic distribution patterns.

**Figure 1.**
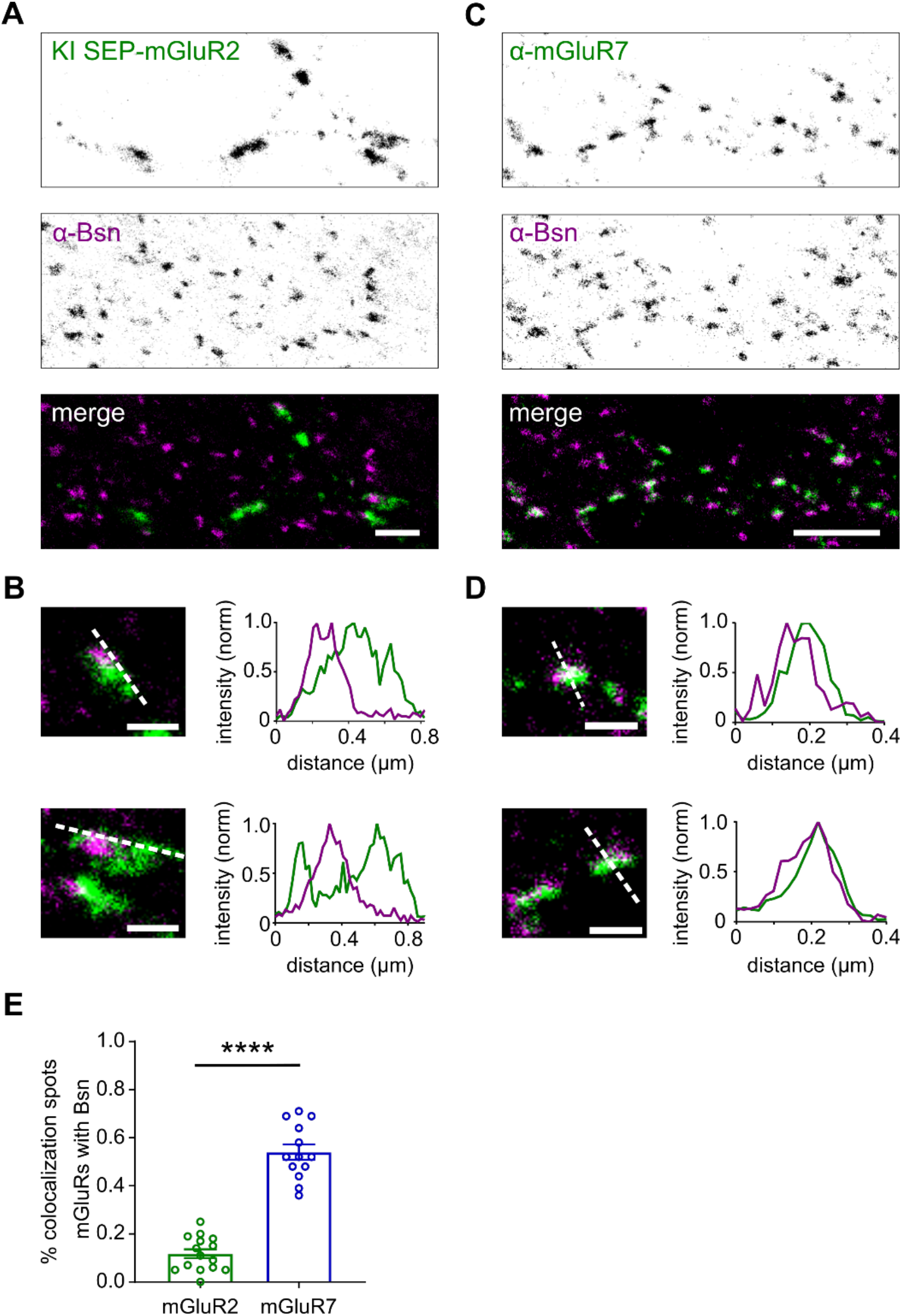
Subsynaptic distribution of presynaptic mGluRs. **(A)** gSTED image of SEP-mGluR2 CRISPR/Cas9 knock-in neuron co-stained with anti-Bassoon (STAR635P) (Bsn). Note that due to the low endogenous expression level of mGluR2, SEP signal was enhanced with anti-GFP (STAR580) staining. Scale bar, 2 µm. **(B)** Example images and intensity profiles of individual mGluR2 positive synapses from (A). Scale bar, 500 nm. **(C)** gSTED image of neuron co-stained with anti-mGluR7 (STAR580) and anti-Bsn (STAR635P). Scale bar, 2 µm. **(D)** Example images and intensity profiles of individual mGluR7-positive synapses from (C). Scale bar: 500 nm. **(E)** Quantification of co-localization between presynaptic mGluRs and Bsn. Unpaired *t-*test, *** *P < 0*.*001*.

**Figure S1.**
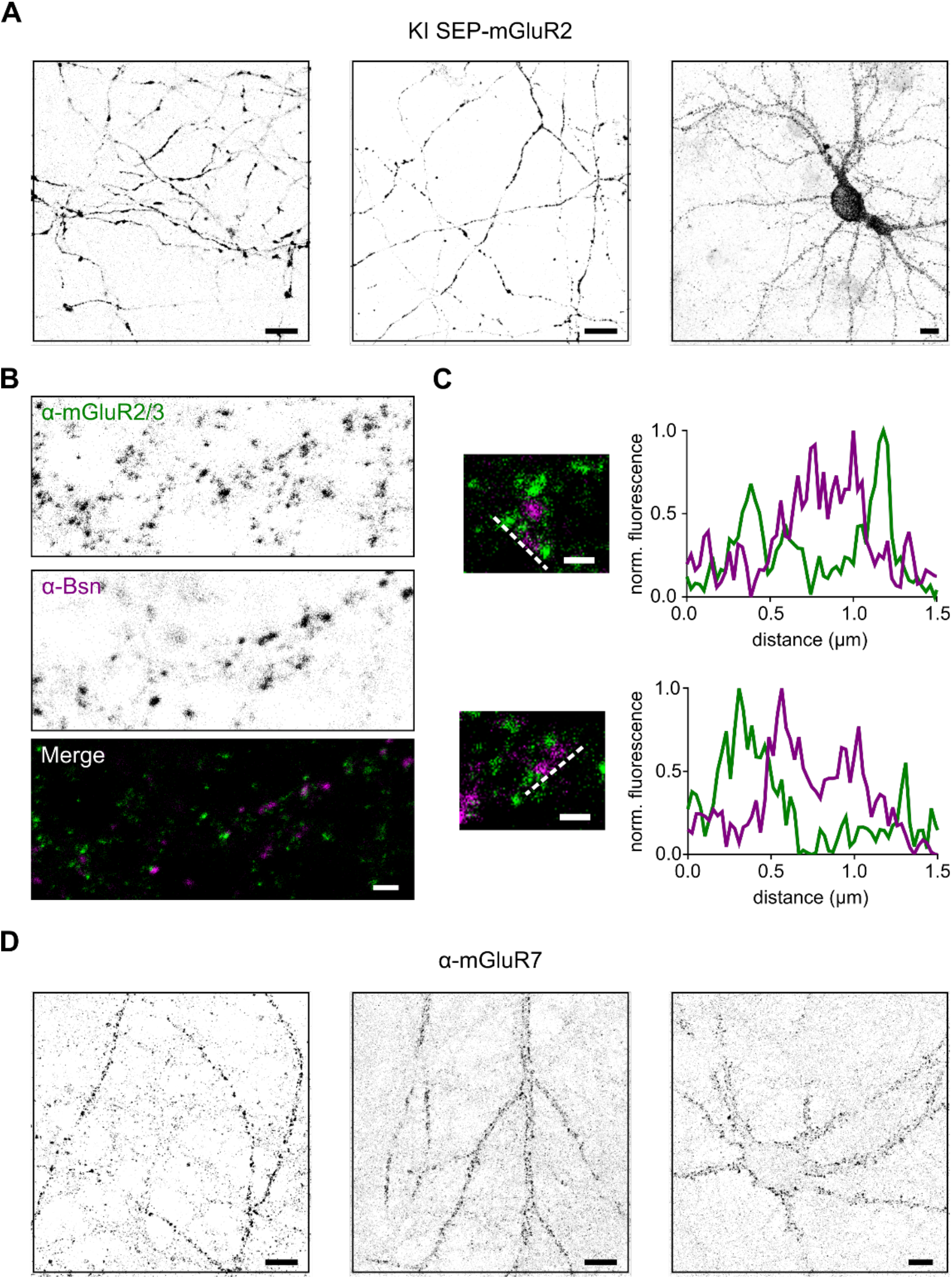
(Related to Figure 1): Distribution of presynaptic mGluRs. **(A)** Example confocal images of SEP-mGluR2 knock-in neurons. SEP signal was enhanced with anti-GFP (STAR580) staining. Scale bar 10 µm. **(B)** gSTED image of neuron co-stained with anti-mGluR2/3 (STAR580) and anti-Bassoon (STAR635P) (Bsn). Scale bar, 2 µm. **(C)** Example images and intensity profiles of individual mGluR2/3 positive synapses from **(B)**. Scale bar, 500 nm. **(D)** Example confocal images of neurons stained with anti-mGluR7 (STAR580). Scale bar, 10 µm.

### Differential stability of mGluR2 and mGluR7 at presynaptic boutons

To test if the observed receptor distributions reflect differences in surface mobility in the axonal membrane, we expressed SEP-tagged mGluR2 and mGluR7 to visualize surface-expressed receptors in live cells and performed fluorescence recovery after photobleaching (FRAP) experiments. Importantly, we found that expressed receptors were efficiently targeted to axons and their localization was consistent with the observed endogenous distributions. SEP-mGluR7 was enriched in presynaptic boutons, while SEP-mGluR2 expression was more diffuse throughout the axon (Fig. S2A-C). Additionally, we verified that expression levels of SEP-tagged mGluRs are comparable (on average ∼ 50% increase) with endogenous levels of mGluR2 and mGluR7 (Fig. S2D, E) We photobleached the fluorescence in small regions overlapping with presynaptic boutons and monitored the recovery of fluorescence over time. Strikingly, the recovery of fluorescence was much more rapid and pronounced for SEP-mGluR2 than for SEP-mGluR7 (Fig. 2A, B). Indeed, quantification of the fluorescence recovery curves showed that the mobile fraction (SEP-mGluR2: 0.60 ± 0.04, SEP-mGluR7: 0.29 ± 0.03, *P<*0.0005, unpaired *t*-test; Fig. 2D) and the recovery time (τ SEP-mGluR2 15.0 ± 1.8 s, SEP-mGluR7 23.5 ± 2.3 s, *P<*0.05, unpaired *t*-test; Fig. 2C) of SEP-mGluR2 were significantly higher than observed for SEP-mGluR7. Thus, these results indicate that mGluR2 is highly mobile in axons, while mGluR7 is immobilized at presynaptic sites and displays minor exchange between synapses.

**Figure 2.**
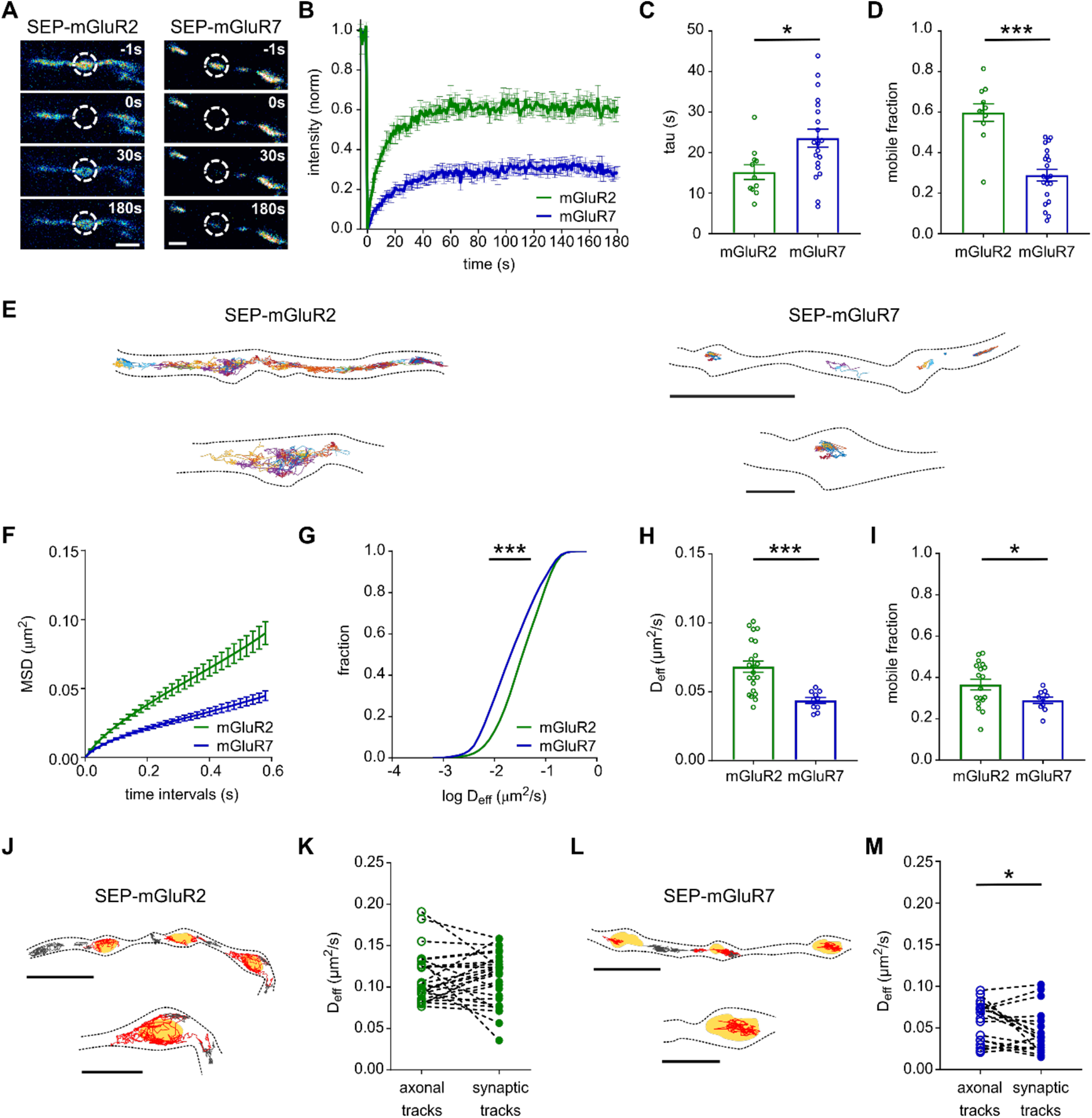
Distinct surface diffusion behavior of mGluR2 and mGluR7. **(A)** Example images from a FRAP time series in neurons expressing SEP-mGluR2 and SEP-mGluR7. The dotted circles indicate the bleached boutons. Scale bar, 1 µm. **(B)** Normalized fluorescence recovery of SEP-mGluR2 and SEP-mGluR7 (n = 11 boutons for SEP-mGluR2, 21 boutons for SEP-mGluR7 from 2 independent experiments). **(C and D)** Quantification of tau of fluorescence recovery (C) and the mobile fraction (D) of SEP-tagged mGluRs. Unpaired *t-*test, *P<0*.*05*, *** *P<0*.*0005*. Error bars represent SEM. **(E)** Example trajectories of SEP-mGluR2 and SEP-mGluR7. Trajectories are displayed with random colors. Outlines of cells are based on the TIRF image of SEP signal. Scale bar, 5 µm; zooms, 1 µm. **(F)** Average mean square displacement (MSD) plot of SEP-mGluR2 and SEP-mGluR7 (n = 22 fields of view for mGluR2, 10 fields of view form mGluR7 from 3 independent experiments) **(G)** Frequency distribution of instantaneous diffusion coefficient (D_eff_) of SEP-mGluR2 and SEP-mGluR7 (n = 22,821 trajectories for SEP-mGluR2, 5,161 trajectories for SEP-mGluR7). Kolmogorov-Smirnov test; *** *P<0*.*0001*. **(H and I)** Quantification of the average instantaneous diffusion coefficient (D_eff_) (H) and the mobile fraction (I) of SEP-tagged mGluRs (n = 22 fields of view for mGluR2, 10 fields of view for mGluR7 from 3 independent experiments). Unpaired *t-*test; *P<0*.*05*, *** *P<0*.*0005*. Error bars represent SEM. **(J)** Trajectories of SEP-mGluR2 plotted on top of the mask marking the presynaptic bouton. Red tracks - synaptic tracks, grey tracks - axonal tracks, yellow areas - bouton mask based on Syp-mCherry signal. Scale bar, 2 µm; zooms, 1 µm. **(K)** Quantification of the instantaneous diffusion coefficient (D_eff_) of axonal and synaptic tracks of SEP-mGluR2 (n = 27 fields of view from 2 independent experiments). **(L)** Trajectories of SEP-mGluR7 plotted on top of the mask marking the presynaptic bouton, as in (J). **(M)** Quantification of instantaneous diffusion coefficient (D_eff_) of axonal and synaptic tracks of SEP-mGluR7 (n = 18 fields of view from 5 independent experiments). Paired *t-*test, * *P<0*.*05*.

**Figure S2.**
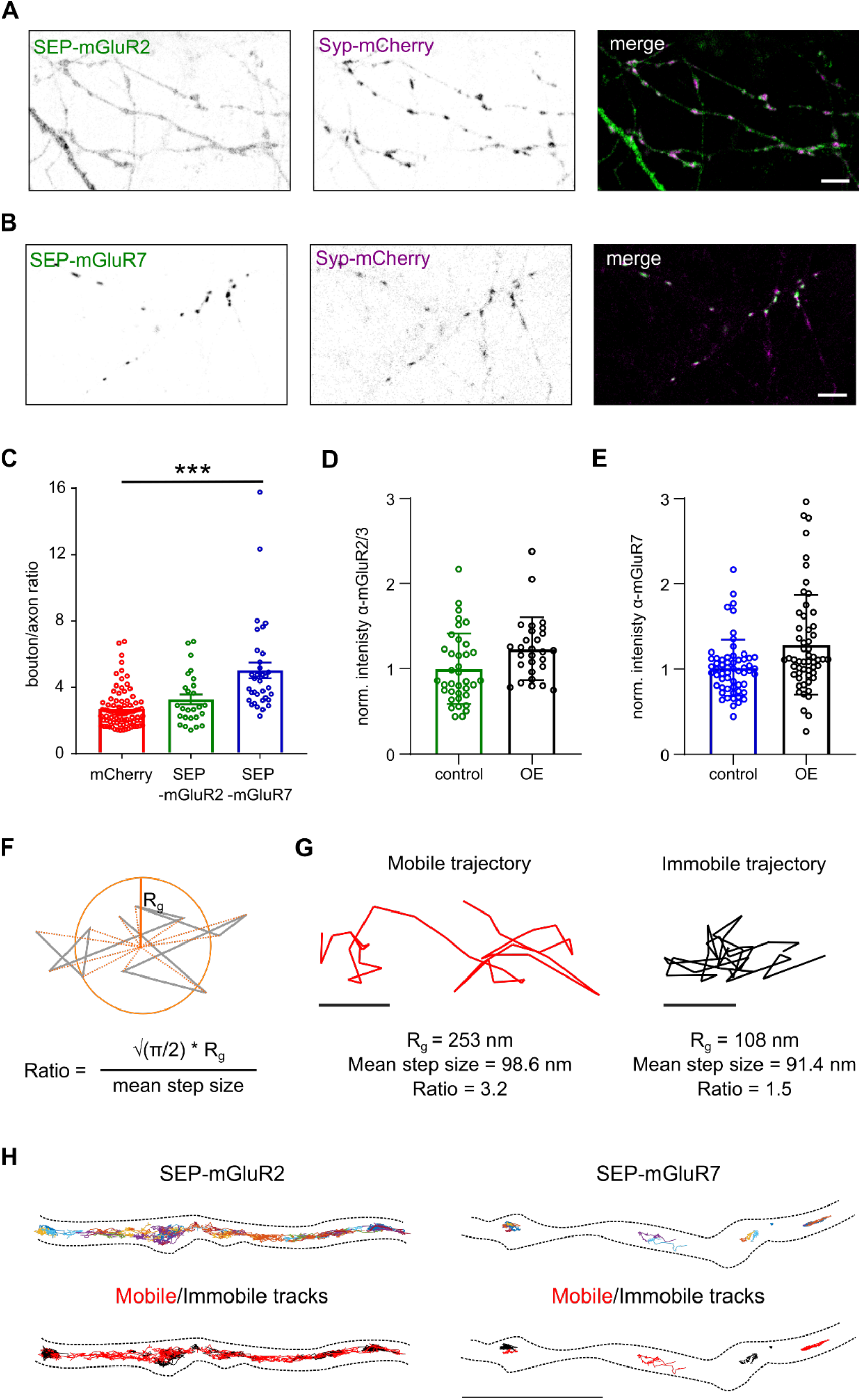
(Related to Figure 2): Targeting of SEP-tagged mGluRs and quantification of mobile and immobile fraction in single-molecule tracking experiments. **(A – B)** An example confocal image of neurons expressing marker of presynaptic boutons Synaptophysin1-mCherry (Syp-mCherry) and SEP-mGluR2 **(A)** or SEP-mGluR7 **(B)**. Scale bar, 5 µm. **(C)** Quantification of ratios of fluorescence intensity in bouton over axon (n = 84 boutons for mCherry, 26 boutons for SEP-mGluR2, 34 boutons for SEP-mGluR7 from 2 independent experiments). The apparent increase in the bouton/axon ratio of cytosolic mCherry likely results from larger bouton volume compared to the axon. One-way ANOVA followed by Dunnett’s multiple comparisons test, *** *P<0*.*001*. **(D - E)** Quantification of overexpression (OE) levels of SEP-tagged presynaptic mGluRs compare to endogenous levels of mGluR2 **(D)** (n = 37 boutons for control, 28 boutons for SEP-mGluR2 overexpression, from 2 independent experiments) and mGluR7 **(E)** (n = 59 boutons for control, 56 boutons for SEP-mGluR7 overexpression, from 2 independent experiments) **(F)** Geometry criterion of mobile/immobile classification of trajectories in single-molecule experiments is based on the ratio between the radius of gyration (Rg) and the mean step size. For an immobile trajectory this ratio is expected to approach 1, while for diffusion particles this ratio is >1. We experimentally validated that the ratio of 2.11 is a valid criterion to differentiate between immobile and mobile tracks. Grey – theoretical trajectory, orange – radius of gyration. **(G)** Examples of mobile and immobile trajectories of mGluR2. Scale bar, 200 nm. **(H)** Trajectory maps of SEP-mGluR2 and SEP-mGluR7 from Figure 2E colored coded for mobile and immobile tracks. Red – mobile trajectories, black – immobile trajectories. Scale bar, 5 µm.

### Single-molecule tracking reveals differences in diffusional behavior of mGluR2 and mGluR7

To resolve the dynamics of mGluR2 and mGluR7 at high spatial resolution and to investigate whether the diffusional behavior of these receptors is heterogeneous within axons, we next performed live-cell single-molecule tracking experiments using universal point accumulation in nanoscale topography (uPAINT) (Giannone *et al*, 2010). SEP-tagged receptors were labeled with anti-GFP nanobodies conjugated to ATTO-647N at low concentrations, which allowed to reliably detect, localize, and track single receptors over time for up to several seconds. The acquired receptor tracks were then compiled into trajectory maps revealing the spatial distribution of receptor motion. These maps were consistent with the receptor distribution patterns as resolved with gSTED imaging. SEP-mGluR2 seemed to rapidly diffuse throughout the axon and synaptic boutons, while SEP-mGluR7 motion was limited and highly confined within synaptic boutons with only a few molecules occasionally diffusing along the axon shaft (Fig. 2E). The mean squared displacement (MSD) vs. elapsed time curves (Fig. 2F) display a sublinear relationship for both receptor types indicating that the majority of these receptors undergo anomalous diffusion. The instantaneous diffusion coefficients (D_eff_) for both receptors was estimated by fitting the slope through the four initial points of the MSD curves. Histograms of D_eff_ estimated from individual trajectories (Fig. 2G) and the average D_eff_ per field of view (Fig. 2H) revealed a significantly higher diffusion coefficient for SEP-mGluR2 than for SEP-mGluR7 (D_eff_ SEP-mGluR2: 0.068 ± 0.004 µm^2^/s, SEP-mGluR7: 0.044 ± 0.002 µm^2^/s, *P<*0.0005, unpaired *t*-test), further indicating that mGluR2 diffuses much more rapidly in the axonal membrane than mGluR7. In addition, we classified the receptors’ diffusional states as either mobile or immobile in a manner independent of MSD-based diffusion coefficient estimation, i.e. by determining the ratio between the radius of gyration and the mean displacement per time step of individual trajectories (Fig. S2F, G) (Golan & Sherman, 2017). Using this approach, we found that SEP-mGluR2 showed a higher fraction of mobile tracks than SEP-mGluR7 (mobile fraction SEP-mGluR2: 0.37 ± 0.03, SEP-mGluR7: 0.29 ± 0.02, *P<*0.05, unpaired *t*-test; Fig. 2I, S2H) further confirming that in axons, mGluR2 is overall more mobile than mGluR7.

To determine whether the surface mobility of these receptors is differentially regulated at synaptic sites, we co-expressed SEP-tagged mGluRs together with a marker of presynaptic boutons, Synaptophysin1 (Syp1) fused to mCherry. Based on epifluorescence images of Syp1-mCherry, we created a mask of presynaptic boutons and compared the D_eff_ of receptors diffusing inside or outside synapses (Fig. 2J, L). The diffusion coefficient of SEP-mGluR2 within presynaptic boutons and along axons did not differ significantly (D_eff_ axonal tracks: 0.113 ± 0.006 µm^2^/s, D_eff_ synaptic tracks: 0.110 ± 0.006 µm^2^/s, *P>*0.05, paired *t*-test; Fig. 2J, K), suggesting that mGluR2 diffusion is not hindered at synaptic sites. Comparing the diffusion coefficients of axonal SEP-mGluR7 tracks with synaptic tracks showed that at a subset of synapses the mobility of SEP-mGluR7 is considerably lower inside boutons (D_eff_ axonal tracks: 0.058 ± 0.006 µm^2^/s, D_eff_ synaptic tracks: 0.045 ± 0.006 µm^2^/s, *P<*0.05, paired *t*-test; Fig. 2L, M), suggesting that mobility of mGluR7 is regulated specifically at the presynaptic sites. Taken together, the FRAP and single-molecule tracking data indicate a striking difference in the dynamic behavior of presynaptic mGluRs. mGluR2 diffuses seemingly unhindered throughout the axon, while mGluR7 is largely immobilized, preferentially at presynaptic active zones.

### The intracellular domain of mGluR2 regulates receptor mobility

To gain insight into the structural mechanisms that control the dynamics of presynaptic mGluRs and to explain the distinct diffusional properties of mGluR2 and mGluR7, we next sought to identify the receptor domains that are involved in controlling mGluR mobility. mGluRs consist of three regions: the intracellular domain (ICD) containing a PDZ binding motif, the prototypical seven-helix transmembrane domain (TMD) involved in G-protein coupling, and the large extracellular domain (ECD) that includes the ligand-binding site (Niswender & Conn, 2010). First, to unravel which segment of mGluR2 regulates its mobility, we created three chimeric receptors of mGluR2 by exchanging the ICD, TMD, or ECD domains of mGluR2 with the corresponding domains of mGluR7 to maintain the overall structure of the receptor. All SEP-tagged chimeric mGluR2 variants were targeted to the axon, similar to wild-type mGluR2, indicating that axonal targeting and surface expression were not altered by replacing these domains (Fig. S3A). Moreover, single-molecule tracking showed that all chimeric mGluR2 variants displayed rapid diffusion throughout the axon and presynaptic boutons, similar to wild-type mGluR2 (Fig. 3A). Interestingly though, the mGluR2 chimera containing the ICD of mGluR7 revealed a significantly higher diffusion coefficient compared to wild-type mGluR2 (D_eff_ SEP-mGluR2-ICD7: 0.082 ± 0.003 µm^2^/s, SEP-mGluR2: 0.065 ± 0.003 µm^2^/s, *P<*0.005, one-way ANOVA), while exchanging the TMD or ECD did not affect the diffusion kinetics of mGluR2 (D_eff_ SEP-mGluR2-TMD7: 0.063 ± 0.004 µm^2^/s; SEP-mGluR2-ECD7: 0.067 ± 0.003 µm^2^/s, *P*>0.05, one-way ANOVA; Fig. 3B). Thus, comparing the diffusional behavior of this set of chimeric mGluR2 variants indicates that intracellular interactions mediate mGluR2 mobility in axons.

**Figure 3.**
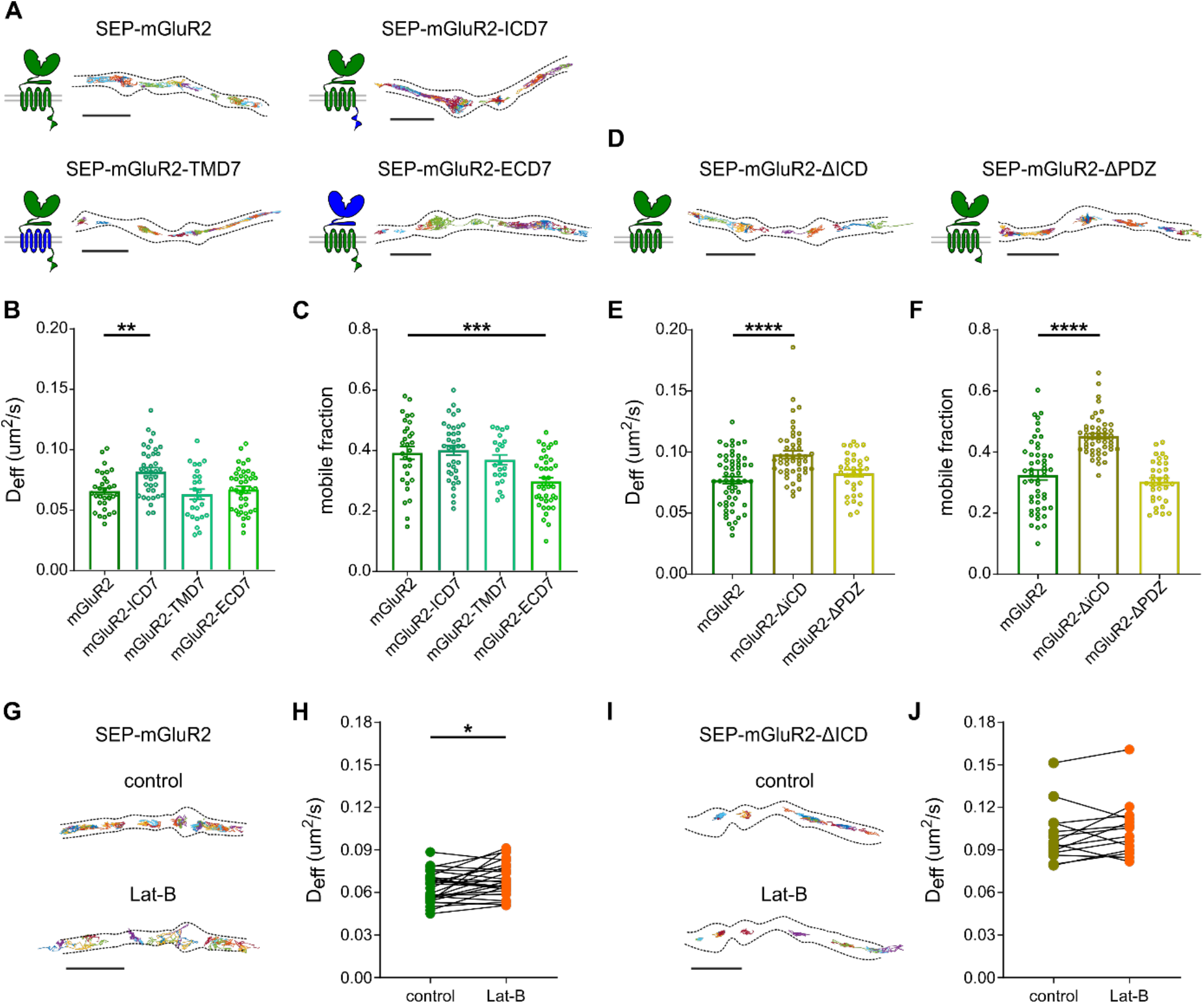
Intracellular interactions regulate the mobility of presynaptic mGluR2. **(A)** Schematic diagrams and example trajectories of wild-type and chimeric variants of mGluR2 (green) with the ICD, TMD, and ECD exchanged with the corresponding mGluR7 domains (blue). Scale bar, 2 µm. **(B and C)** Quantification of average diffusion coefficient (D_eff_) (B) and the mobile fraction (C) of SEP-tagged chimeric mGluR2 variants (n = 30 - 40 fields of view from 4 - 5 independent experiments). **(D)** Schematic diagrams and example trajectories of C-terminal deletion variants of mGluR2. Scale bar, 2 µm. **(E and F)** Quantification of average diffusion coefficient (D_eff_) (E) and the mobile fraction (F) of SEP-tagged C-terminal deletion variants of mGluR2 (n = 34 – 53 fields of view from 3 independent experiments). One-way ANOVA followed by Dunnett’s multiple comparison test; ** *P<0*.*005*, *** *P<0*.*0005*. **(G)** Example trajectories of SEP-mGluR2 before and after incubation with 5 µM latrunculin B (Lat-B). Scale bar, 2 µm. **(H)** Quantification of diffusion coefficient (D_eff_) of SEP-mGluR2 before and after incubation with Lat-B (n = 27 fields of view, from 3 independent experiments). Paired *t-*test, * *P<0*.*05*. **(I)** Example trajectories of SEP-mGluR2-ΔICD before and after incubation with Lat-B. Scale bar, 2 µm. **(J)** Quantification of diffusion coefficient (D_eff_) of SEP-mGluR2-ΔICD before and after incubation with Lat-B (n = 14, from 2 independent experiments). Error bars represent SEM. All trajectories are displayed with random colors. Outlines of cells are based on the TIRF image of SEP signal.

To further investigate whether intracellular interactions regulate mGluR2 dynamic, we created two deletion variants of mGluR2 by removing the entire ICD or only the distal PDZ binding motif. SEP-tagged C-terminal deletion mGluR2 variants were targeted to the axon, similar to the wild-type receptor (Fig. S3B). Also, trajectory maps revealed their rapid diffusion throughout the axon and presynaptic boutons (Fig. 3D). Interestingly, deletion of the PDZ binding motif did not alter the diffusion behavior of mGluR2, however, removal of the entire ICD increased the diffusion coefficient (D_eff_ SEP-mGluR2-ΔICD: 0.098 ± 0.003 µm^2^/s, SEP-mGluR2-ΔPDZ: 0.082 ± 0.003 µm^2^/s, SEP SEP-mGluR2: 0.077 ± 0.003 µm^2^/s, *P<*0.0001, one-way ANOVA; Fig. 3E) and the mobile fraction of mGluR2 (mobile fraction SEP-mGluR2-ΔICD: 0.45 ± 0.01, SEP-mGluR2-ΔPDZ: 0.30 ± 0.01, SEP-mGluR7: 0.32 ± 0.02, *P*<0.0001, one-way ANOVA; Fig. 3F) confirming that the ICD of mGluR2 is involved in controlling receptor mobility but, this regulation is not mediated by interactions via PDZ-mediated interactions.

Little is known about mGluR2 C-tail-mediated interactions, but we reasoned that direct or indirect interactions with the actin cytoskeleton, which have an important role in controlling membrane organization in axons (He *et al*, 2016; Zhou *et al*, 2019), could influence mGluR2 diffusion. Disruption of the actin cytoskeleton with latrunculin-B (Lat-B) increased the diffusion coefficient of wild-type mGluR2 (D_eff_ control: 0.063 ± 0.002 µm^2^/s, Lat-B: 0.069 ± 0.002 µm^2^/s, *P<*0.05, paired t-test; Fig. 3G, H) but did not cause a further increase in the diffusion coefficient of mGluR2 lacking the ICD (D_eff_ control: 0.100 ± 0.005 µm^2^/s, Lat-B: 0.104 ± 0.005 µm^2^/s, *P>*0.05, paired t-test; Fig. 3I, J). These results suggest that intracellular interactions with the actin cytoskeleton regulate mGluR2 mobility.

**Figure S3.**
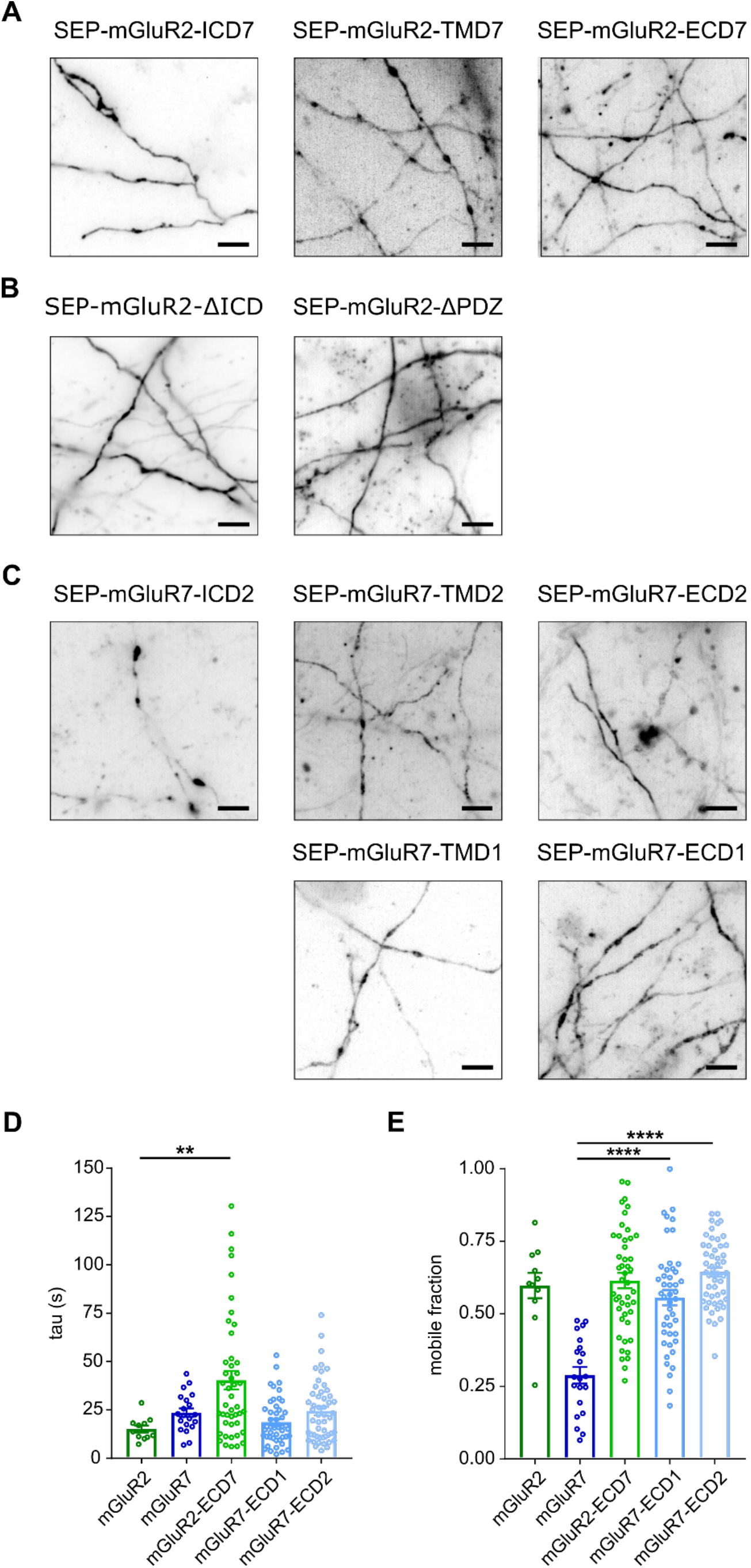
(Related to Figure 3 and 4): Expression of mGluRs variants and FRAP experiments of extracellular chimeric receptors. **(A)** Example images of neurons expressing SEP-tagged chimeric mGluR2 variants. Scale bar, 5 µm. **(B)** Example images of neurons expressing SEP-tagged C-terminal deletion mGluR2 variants Scale bar, 5 µm. **(C)** Example images of neurons expressing SEP-tagged chimeric mGluR7 variants. Scale bar, 5 µm. **(D and E)** Quantification of half time of fluorescence recovery **(D)** and mobile fraction **(E)** from FRAP experiments of SEP-tagged extracellular chimeric variants of mGluR2 and mGluR7 (n = 10 - 45 boutons from 2 - 3 independent experiments). Kruskal-Wallis ANOVA in **(D)**; one-way ANOVA in **(E)** followed by Dunnett’s multiple comparisons test ** *P<0*.*05*, **** *P<0*.*0005*. Error bars represent SEM.

### mGluR7 immobilization at presynaptic active zones is controlled by extracellular domain

While mGluR2 rapidly diffuses through the axon, we found that mGluR7 is stably anchored and concentrated at active zones. Therefore, we decided to further focus on the mechanisms that could underlie the immobilization of mGluR7 at presynaptic sites. To test which region of mGluR7 is involved in the immobilization of mGluR7 at the active zone, we generated five chimeric variants of mGluR7 to exchange the ICD, TMD, or ECD of mGluR7 with the corresponding domains of mGluR2 or mGluR1. Because the C-terminal domain of mGluR1 is involved in targeting the receptor to the dendritic compartment we decided to not substitute the ICD of mGluR7 for the ICD of mGluR1 (Francesconi & Duvoisin, 2002). All SEP-tagged chimeric variants of mGluR7 were readily detected in axons, similar to wild-type mGluR7 (Fig. S3C) indicating that these receptors are correctly targeted to the axonal membrane.

In contrast to mGluR2, the exchange of the ICD of mGluR7 did not change the diffusional behavior of the receptor. Trajectory maps obtained from single-molecule tracking showed that diffusion of the SEP-tagged mGluR7 chimera containing the ICD of mGluR2 was still restricted to presynaptic boutons (Fig. 4A) and the diffusion coefficient (D_eff_ SEP-mGluR7-ICD2: 0.043 ± 0.004 µm^2^/s, SEP-mGluR7: 0.039 ± 0.002 µm^2^/s, *P*>0.05, one-way ANOVA; Fig. 4B) and mobile fraction were similar to wild-type SEP-mGluR7 (mobile fraction SEP-mGluR7-ICD2: 0.18 ± 0.03, SEP-mGluR7: 0.21 ± 0.02, *P*>0.05, one-way ANOVA; Fig. 4C), suggesting that intracellular interactions do not contribute to mGluR7 immobilization. Diffusion of SEP-tagged TMD chimeric variants of mGluR7 was also mostly restricted to presynaptic boutons (Fig. 4A), although we found that replacing the mGluR7 TMD with the TMD of mGluR2 slightly increased the diffusion coefficient (D_eff_: SEP-mGluR7-TMD2 0.059 ± 0.004 µm^2^/s, *P*<0.05, one-way ANOVA; Fig. 4B) and the mobile fraction (SEP-mGluR7-TMD2 0.29 ± 0.02, *P*<0.05, one-way ANOVA; Fig. 4C). However, the substitution of the mGluR7 TMD with the mGluR1 TMD did not alter its diffusional behavior (D_eff_ SEP-mGluR7-TMD1: 0.033 ± 0.003 µm^2^/s, mobile fraction: 0.16 ± 0.02, *P*>0.05, one-way ANOVA; Fig. 4B, C), suggesting that the faster diffusion of the mGluR7 variant containing the TMD of mGluR2 is most likely due to specific properties of the mGluR2 TMD and cannot be attributed to a mGluR7-specific mechanism. Indeed, a previous study reported stronger interactions between transmembrane regions in mGluR2 homodimers compared to other mGluR subtypes (Gutzeit *et al*, 2019).

**Figure 4.**
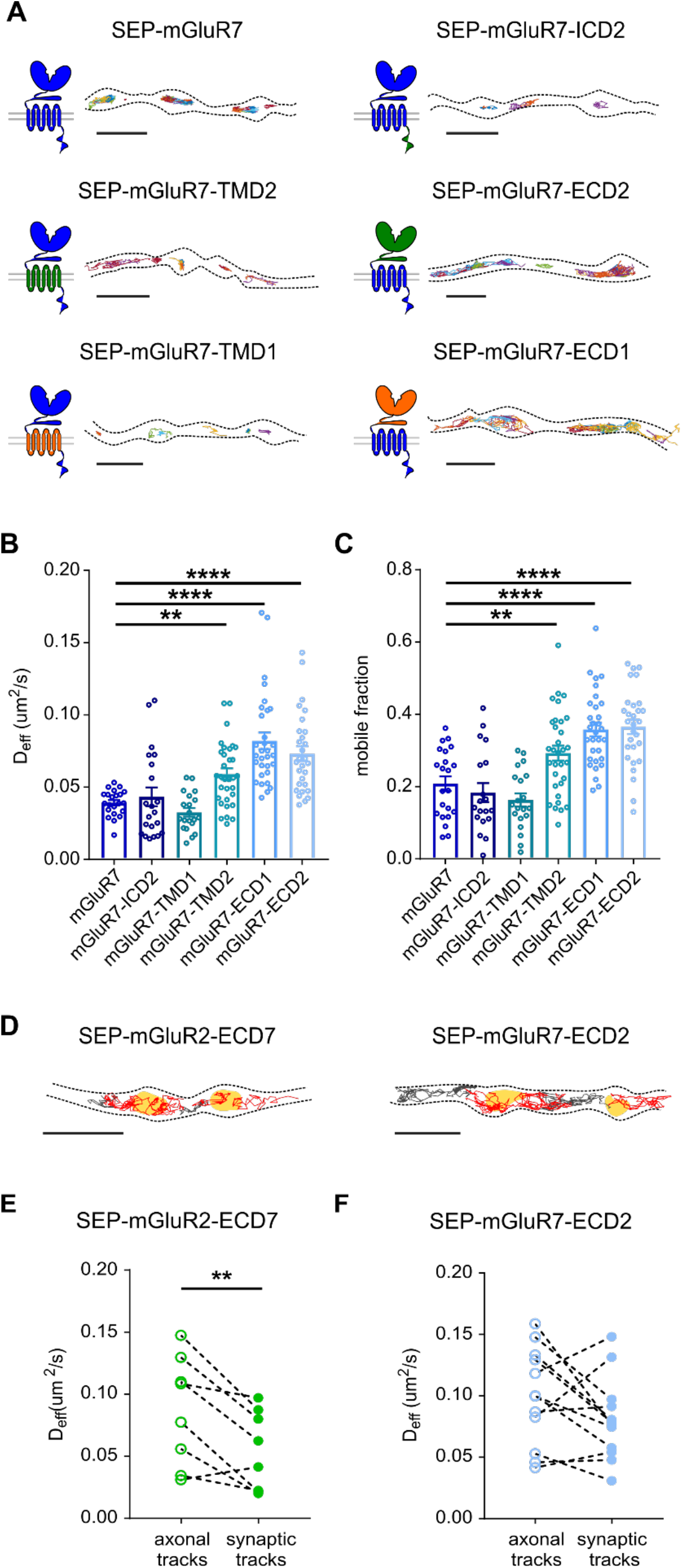
Extracellular domain regulates the mobility of mGluR7. **(A)** Schematic diagrams and example trajectories of wild-type and chimeric variants of mGluR7 (blue) with the ICD, TMD, and ECD exchanged with the corresponding domains from mGluR2 (green) or mGluR1 (orange). Scale bar, 2 µm. **(B and C)** Quantification of average diffusion coefficient (D_eff_) (B) and the mobile fraction (C) of SEP-tagged chimeras of mGluR7 (n = 22 - 32 fields of view from 4 - 5 independent experiments). One-way ANOVA followed by Dunnett’s multiple comparison test; ** *P<0*.*05*, **** *P<0*.*0005*. **(D)** Trajectories of extracellular chimeras SEP-mGluR2-ECD7 and SEP-mGluR7-ECD2 plotted on top of the mask of the presynaptic bouton. Red tracks - synaptic tracks, grey tracks - axonal tracks, yellow areas - bouton mask based on Syp-mCherry signal. Scale bar, 2 µm. **(E and F)** Quantification of diffusion coefficient (D_eff_) of axonal and synaptic tracks of SEP-mGluR2-ECD7 (E) and SEP-mGluR7-ECD2 (F) (n = 8 fields of view for SEP-mGluR2-ECD7, 12 fields of view for SEP-mGluR7-ECD2 from 2 independent experiments). Paired *t-*test, ** *P<0*.*005*. Error bars represent SEM. All trajectories are displayed with random colors. Outlines of cells are based on the TIRF image of SEP signal.

Interestingly, replacing the ECD of mGluR7 drastically altered its diffusional behavior. In contrast to the wild-type receptor, SEP-tagged chimeric mGluR7 variants containing the ECD of mGluR2 or mGluR1 diffused freely throughout the axon and boutons (Fig. 4A) and displayed almost a two-fold increase in diffusion coefficient (D_eff_ SEP-mGluR7-ECD1: 0.082 ± 0.006 µm^2^/s, SEP-mGluR7-ECD2: 0.073 ± 0.005 µm^2^/s, *P<*0.0005, one-way ANOVA; Fig. 4B) and larger mobile fraction compared to wild-type SEP-mGluR7 (SEP-mGluR7-ECD1: 0.36 ± 0.02, SEP-mGluR7-ECD2: 0.37 ± 0.02, SEP-mGluR7: 0.21 ± 0.02, *P*<0.0005, one-way ANOVA; Fig. 4C). Thus, the immobilization of mGluR7 at presynaptic sites likely relies on extracellular interactions with its ECD. To assess if the ECD of mGluR7 is sufficient to immobilize receptors, we replaced the ECD of mGluR2 with the ECD of mGluR7. Indeed, we found a significant decrease in the mobile fraction of the SEP-tagged chimeric mGluR2 variant containing the mGluR7 ECD (SEP-mGluR2-ECD7: 0.30 ± 0.02, SEP-mGluR2: 0.39 ± 0.02, *P<*0.0005, one-way ANOVA; Fig. 3C) supporting the role of the mGluR7 ECD in immobilizing the receptor. To further substantiate these results, we performed FRAP experiments and found a significant increase in fluorescence recovery of SEP-tagged mGluR7 variants with substituted ECDs (Fig. S3E) and slower recovery kinetics of SEP-tagged chimeric mGluR2 with the ECD of mGluR7 (Fig. S3D). These results are in striking agreement with the single-molecule tracking data and confirm the dominant role of the mGluR7 ECD in regulating receptor mobility.

Based on our findings that the localization of mGluR7 is restricted to the active zone and that mGluR7 diffusion is hindered at presynaptic boutons, we hypothesized that the ECD of mGluR7 mediates receptor immobilization specifically at presynaptic sites. To test this, we resolved receptor mobility at synapses by co-expressing ECD chimeric variants of mGluR2 and mGluR7 with Syp1-mCherry (Fig. 4D). Although the mGluR2 chimera containing the ECD of mGluR7 displayed rather high diffusion coefficients in the axonal shaft, the pool of chimeric receptors inside presynaptic boutons showed a significantly lower diffusion coefficient (D_eff_ synaptic tracks: 0.054 ± 0.011 µm^2^/s, axonal tracks: 0.087 ± 0.015 µm^2^/s, *P*<0.005, paired *t*-test; Fig. 4E). Vice versa, replacing the ECD of mGluR7 with the ECD of mGluR2 resulted in a similar diffusion coefficient of axonal and synaptic tracks (D_eff_ synaptic tracks: 0.081 ± 0.01 µm^2^/s, axonal tracks: 0.1 ± 0.01 µm^2^/s, *P*>0.05, paired *t*-test; Fig. 4F) suggesting that the ECD of mGluR7 is indeed sufficient to immobilize receptors at presynaptic sites. Altogether, these results indicate that mGluR7 immobilization at synaptic sites is in large part mediated by extracellular interactions.

### Surface mobility of presynaptic mGluRs is altered by decreased but not increased synaptic activity

Our results so far suggest that, under resting conditions, the diffusional properties of presynaptic mGluRs are largely controlled by distinct intra- and extracellular interactions. However, ligand-induced activation of GPCRs involves a dramatic change in receptor conformation and has been shown to change the oligomerization and diffusion behavior of various GPCRs, including mGluRs, in non-neuronal cells (Yanagawa *et al*, 2018; Calebiro *et al*, 2013; Kasai & Kusumi, 2014; Sungkaworn *et al*, 2017). To test whether receptor activation alters the diffusion of presynaptic mGluRs in neurons, we performed single-molecule tracking of mGluR2 and mGluR7 before and after stimulation with their specific agonists. We found that activation of SEP-mGluR2 with the selective group II mGluR agonist LY379268 (LY) did not change the distribution of receptor trajectories (Fig. 5A) or diffusion coefficients (D_eff_ control: 0.06 ± 0.003 µm^2^/s, LY: 0.058 ± 0.004 µm^2^/s, *P*>0.05, paired *t*-test; Fig. 5B). Similarly, direct activation of mGluR7 with the potent group III mGluR agonist L-AP4 also did not change the diffusional behavior of SEP-mGluR7 (D_eff_ control: 0.044 ± 0.002 µm^2^/s, L-AP4: 0.045 ± 0.003 µm^2^/s; *P*>0.05, paired *t*-test; Fig. 5C, D). Thus, these experiments indicate that in neurons, the dynamics of presynaptic mGluRs are not modulated by agonist-stimulated receptor activation.

**Figure 5.**
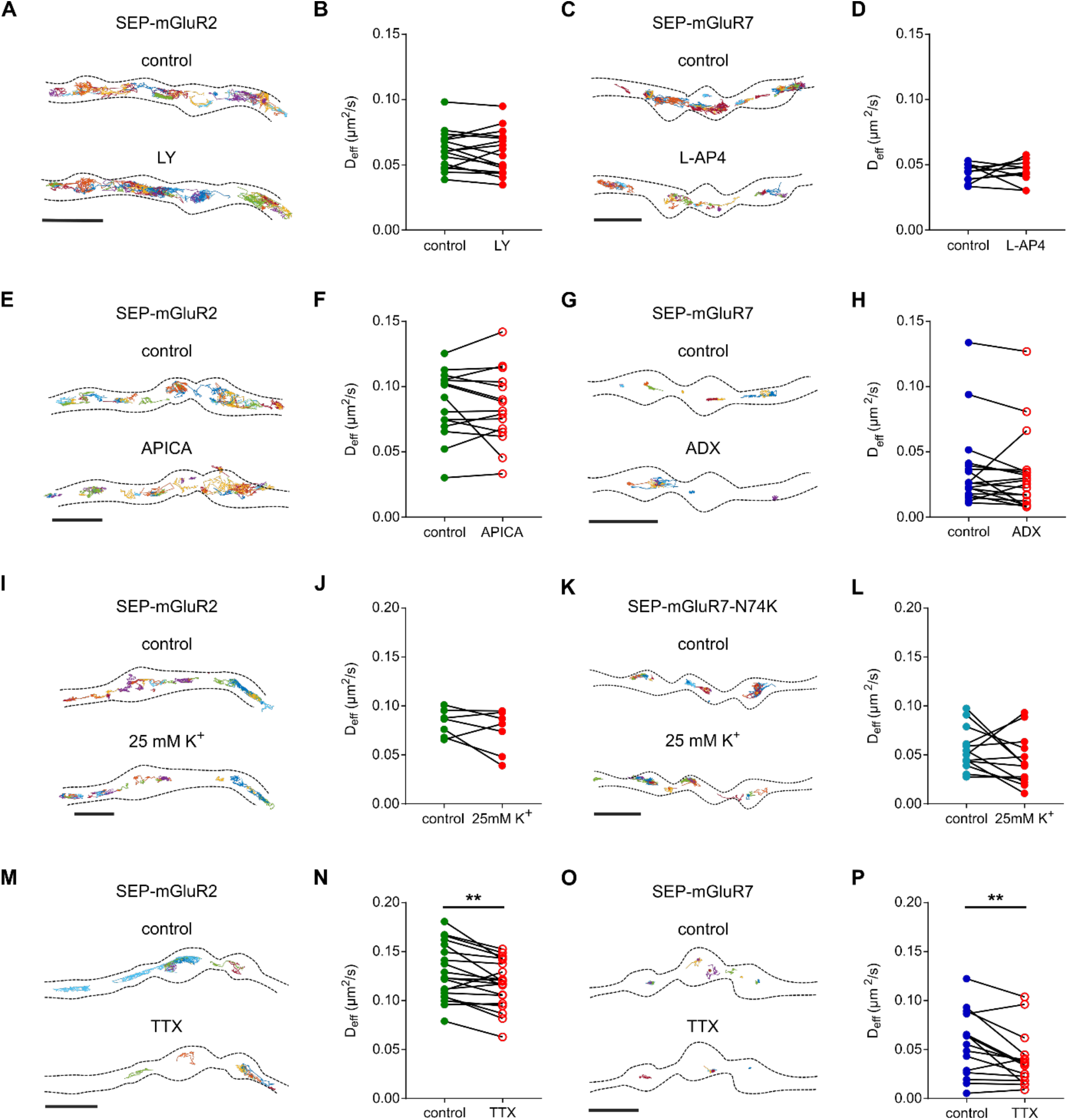
Lateral diffusion of presynaptic mGluRs is not regulated by activity. **(A)** Example trajectories of SEP-mGluR2 before and after incubation with 100 µM LY. Scale bar, 2 µm. **(B)** Quantification of diffusion coefficient (D_eff_) of SEP-mGluR2 before and after incubation with LY (n = 17 fields of view from 2 independent experiments). **(C)** Example trajectories of SEP-mGluR7 before and after incubation with 500 µM L-AP4. Scale bar, 2 µm. **(D)** Quantification of diffusion coefficient (D_eff_) of SEP-mGluR7 before and after incubation with L-AP4 (n = 10 fields of view from 2 independent experiments). **(E)** Example trajectories of SEP-mGluR2 before and after incubation with 100 µM APICA. Scale bar, 2 µm. **(F)** Quantification of diffusion coefficient (D_eff_) of SEP-mGluR2 before and after incubation with APICA (n = 15 fields of view from 3 independent experiments). **(G)** Example trajectories of SEP-mGluR7 before and after incubation with 10 µM ADX. Scale bar, 2 µm. **(H)** Quantification of diffusion coefficient (D_eff_) of SEP-mGluR7 before and after incubation with ADX (n = 18, from 4 independent experiments). **(I)** Example tracks of SEP-mGluR2 before and after incubation with 25 mM K^+^. Scale bar, 2 µm. **(J)** Quantification of diffusion coefficient (D_eff_) of SEP-mGluR2 before and after incubation with 25 mM K^+^ (n = 7 fields of view from 2 independent experiments). **(K)** Example tracks of SEP-mGluR7-N74K before and after incubation with 25 mM K^+^. Scale bar, 2 µm. **(L)** Quantification of diffusion coefficient (D_eff_) of SEP-mGluR7-N74K before and after incubation and with 25 mM K^+^ (n = 13 fields of view from 2 independent experiments). **(M)** Example trajectories of SEP-mGluR2 before and after incubation with 1 µM TTX. Scale bar, 2 µm. **(N)** Quantification of diffusion coefficient (D_eff_) of SEP-mGluR2 before and after incubation with TTX (n = 20 from 3 independent experiments). **(O)** Example trajectories of SEP-mGluR7 before and after incubation with 1 µM TTX. Scale bar, 2 µm. **(P)** Quantification of diffusion coefficient (D_eff_) of SEP-mGluR7 before and after incubation with TTX (n = 15 from 3 independent experiments). All trajectories are displayed with random colors. Outlines of cells are based on the TIRF image of SEP signal. Paired *t-*test, ** *P<0*.*005*.

Next, we tested whether inhibition of presynaptic mGluRs influences their mobility. We found that inhibition of SEP-mGluR2 with the group II mGluR antagonist APICA did not alter the diffusion rate of mGluR2 (D_eff_ control: 0.087 ± 0.007 µm^2^/s, APICA: 0.084 ± 0.007 µm^2^/s, *P*>0.05, paired *t*-test; Fig. 5E, F). Similarly, inhibition of mGluR7 activity with the negative allosteric modulator ADX71743 (ADX) did not change the distribution of trajectories (Fig. 5G) or the diffusion coefficient of the receptor (D_eff_ control: 0.036 ± 0.007 µm^2^/s, ADX: 0.033 ± 0.007 µm^2^/s, *P*>0.05, paired *t*-test; Fig. 5H). Altogether, these results suggest that the diffusion of presynaptic mGluRs is not impacted by changes in the activation state of the receptor.

Global changes in neuronal activity could alter receptor mobility, either directly by receptor stimulation by their endogenous ligand glutamate, or perhaps indirectly through structural changes in synapse organization. To test this, we next determined whether strong synaptic stimulation by application of the potassium channel blocker 4-AP together with the glutamate reuptake blocker TBOA, to acutely increase synaptic glutamate levels, changed receptor diffusion. However, we did not find a significant effect of synaptic stimulation on the diffusion coefficient of SEP-mGluR2 (D_eff_ control: 0.085 ± 0.011 µm^2^/s, 4-AP + TBOA: 0.069 ± 0.009 µm^2^/s, *P*>0.05, paired t-test; Fig. S4A, B). Additionally, even under strong depolarizing conditions (25 mM K^+^, 5 - 10 min), the diffusion coefficient of SEP-mGluR2 remained unaltered (D_eff_ control: 0.082 ± 0.005 µm^2^/s, 25 mM K^+^: 0.074 ± 0.008 µm^2^/s, *P*>0.05, paired *t*-test; Fig. 5I, J). We found similar results for SEP-mGluR7 (data not shown). However, since the affinity of mGluR7 for glutamate is very low, in the range of 0.5 - 1 mM (Schoepp *et al*, 1999), we reasoned that the unaltered diffusion of mGluR7 during synaptic stimulation could be due to the incomplete activation of the receptor. Therefore, we analyzed the mobility of an mGluR7 mutant with a two-fold increased affinity for glutamate (mGluR7-N74K) (Kang *et al*, 2015) during strong depolarization. Importantly, we found that the diffusion rate of SEP-mGluR7-N74K was not significantly different from wild-type SEP-mGluR7 under control conditions (D_eff_ SEP-mGluR7-N74K: 0.049 ± 0.005 µm^2^/s, SEP-mGluR7: 0.039 ± 0.002 µm^2^/s, *P*>0.05, unpaired *t*-test; Fig. S4C-E). However, despite having a two-fold higher affinity for glutamate, the diffusion kinetics of SEP-mGluR7-N74K remained unaltered under strong depolarizing conditions (D_eff_ control: 0.056 ± 0.006 µm^2^/s, 25 mM K^+^: 0.044 ± 0.007 µm^2^/s, *P*>0.05, paired *t*-test; Fig. 5K, L). Thus, these single-molecule tracking experiments demonstrate that the lateral diffusion of presynaptic mGluRs on the axonal membrane is not modulated by direct activation with ligands, or an acute increase in neuronal activity.

Furthermore, we determined whether a reduction in neuronal activity modulates the mobility of presynaptic mGluRs. We analyzed the diffusion of mGluRs after acute inhibition of spontaneous action potential by blocking sodium channels with TTX. Interestingly, we found a reduction of SEP-mGluR2 mobility after treatment with TTX (D_eff_ control: 0.129 ± 0.006 µm^2^/s, TTX: 0.117 ± 0.005 µm^2^/s, *P*<0.005, paired *t*-test; Fig. 5M, N). Moreover, inhibition of spontaneous activity also decreased the diffusion coefficient of SEP-mGluR7 (D_eff_ control: 0.055 ± 0.009 µm^2^/s, TTX: 0.040 ± 0.007 µm^2^/s, *P*<0.05, paired *t*-test; Fig. 5M, N). Thus, our data revealed that the mobility of presynaptic mGluRs is reduced under conditions of reduced neuronal activity.

**Figure S4.**
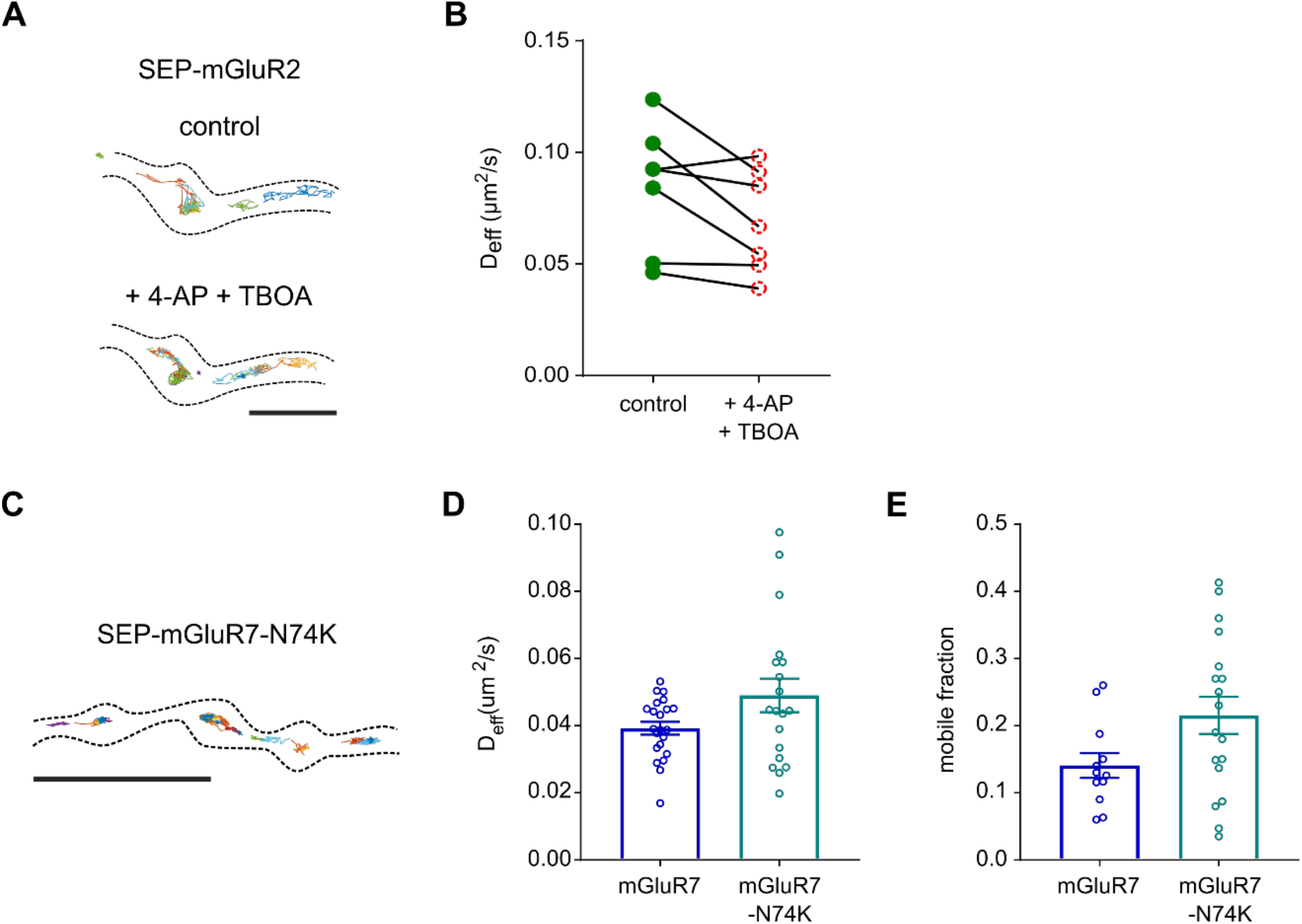
(Related to Figure 5): Mobility of presynaptic mGluR2 does not depend of neuronal activity and high-affinity mutant of mGluR7 displays similar mobility as wild-type receptor. **(A)** Example tracks of SEP-mGluR2 before and after incubation with 200 µM 4-AP and 10 µM TBOA. Scale bar, 2 µm. **(B)** Quantification of diffusion coefficient (D_eff_) of SEP-mGluR2 before and after incubation with 4-AP and TBOA (n = 7 fields of view from 2 independent experiments). **(C)** Example trajectories of SEP-mGluR7-N74K.. Scale bar, 5 µm. **(D and E)** Quantification of average diffusion coefficient (D_eff_) **(D)** and mobile fraction **(E)** of SEP-mGluR7 and mutant SEP-mGluR7-N74K (n = 22 fields of views for SEP-mGluR7, 19 fields of view for SEP-mGluR7-N74K from 2 independent experiments). Trajectories are displayed with random colors. Outlines of cells is based on TIRF image of SEP signal. Error bars represent SEM.

### A computational model of presynaptic mGluR activation reveals that different levels of receptor activation depend on subsynaptic localization

Our data show that mGluR7 is immobilized at the active zone, close to the release site, while mGluR2 is distributed along the axon and synaptic boutons, seemingly excluded from the active zone. Moreover, their localization and dynamics did not change during increased synaptic activity. We hypothesized that these distinct distribution patterns differentially influence the contribution of presynaptic mGluRs to the modulation of synaptic transmission. To test this hypothesis, we investigated a computational model of presynaptic mGluR activation combining the cubic ternary complex activation model (cTCAM) of GPCRs signaling (Fig. 6B) (Kinzer-Ursem & Linderman, 2007) with a model of time-dependent diffusion of glutamate release after a single synaptic vesicle (SV) fusion or multi-vesicle release at different frequencies. To determine the effect of mGluR localization, we compared receptor activation at varying distances (5 nm to 1 µm) from the release site (Fig. 6A). We calibrated the activation model of mGluR2 and mGluR7 by solving cTCAM with different values of the association constant (K_a_), keeping other parameters constant (Table S1), to match the model outputs: the relative number of receptor-ligand complexes (Fig. S5A) and the Gα_GTP_ concentration (Fig. S5B) with previously published EC_50_ values for mGluR2 and mGluR7 (Schoepp *et al*, 1999). Because two out of four liganded receptor states in the cTCAM represent an inactive receptor, we used the G_αGTP_ concentration as a readout of receptor activation to compare responses of mGluRs to different synaptic activity patterns.

**Figure 6.**
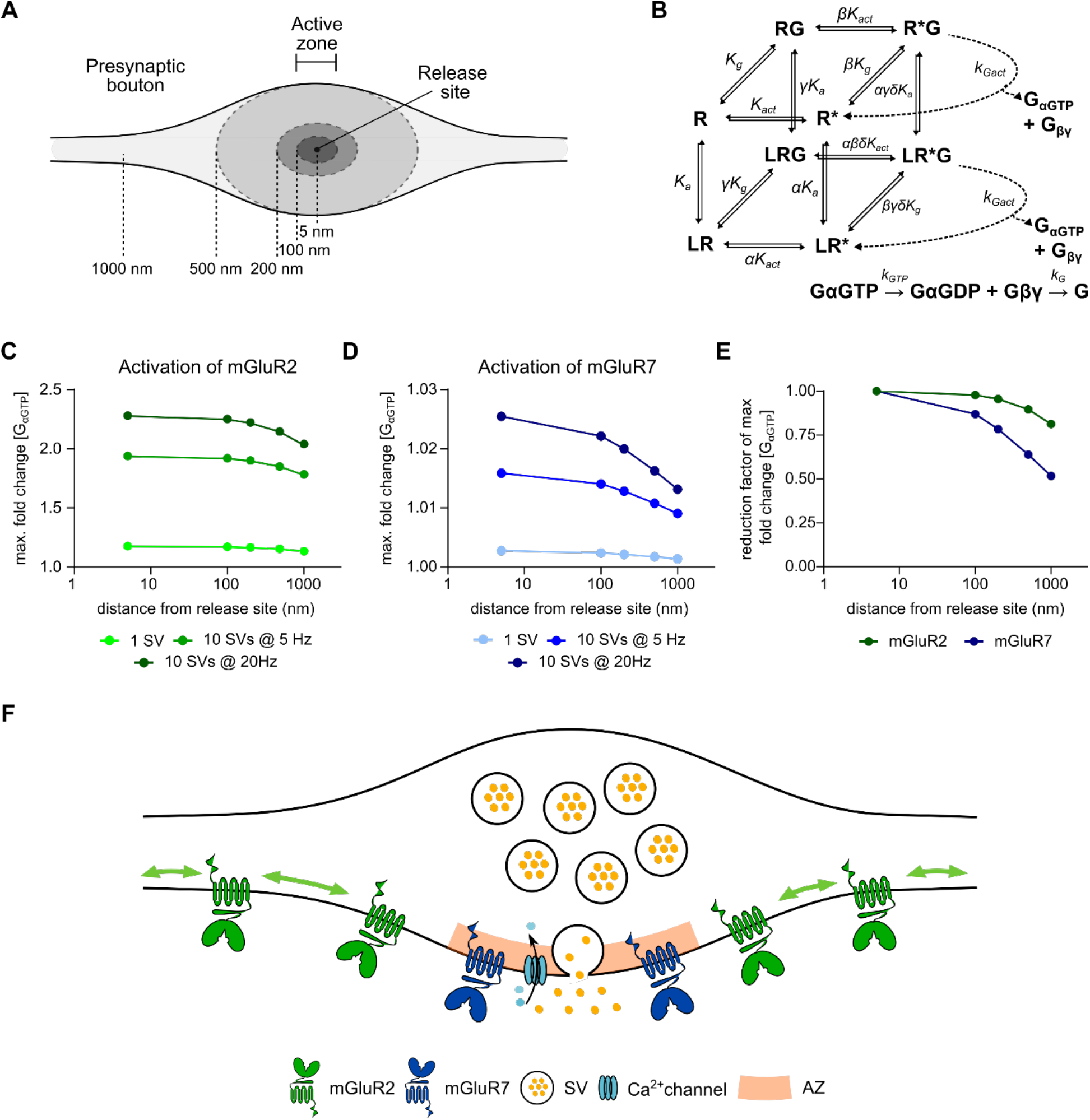
A computational model of mGluRs activation shows that subsynaptic localization of presynaptic mGluRs tunes receptor activation. **(A)** Schematic of presynaptic bouton highlighting subsynaptic localizations used in modeling. **(B)** Kinetics and rate equations described in the cubic ternary complex activation model of presynaptic mGluRs signaling. All parameters used in the model are summarized in Table S1. **(C and D)** Receptor response to glutamate release during different release patterns (1 SV, 10 SVs at 5 Hz, and 10 SVs at 20 Hz) at different distances from release site (5 nm to 1 µm) for mGluR2 (C) and mGluR7 (D). Note that x-axis is on a logarithmic scale. **(E)** The normalized reduction factor of mGluR2 and mGluR7 activation at different distances from the release site after release of 10 SVs at 20Hz. **(F)** A model of subsynaptic distribution and mobility of presynaptic mGluRs. mGluR2 is distributed along the axon and displays high mobility that is modulated by its intracellular interactions. mGluR7 is enriched and immobilized at the active zone. Immobilization of mGluR7 is regulated by its extracellular domain. SV - synaptic vesicle, AZ - the active zone.

The release of glutamate from a single SV, representing release during spontaneous synaptic activity, caused only a slight increase in the activation of mGluR2 when located close to the release site (r = 5 nm) and outside the active zone (r ≥ 100 nm, Fig. 6C, S5C). Release of 10 SVs, corresponding to the size of the readily releasable pool, at low frequency (5 Hz) increased the activity of mGluR2 almost 2-fold inside presynaptic boutons (r ≤ 500 nm; Fig. 6C, S5D). Elevation of the fusion frequency to 20 Hz further increased receptor activation to ∼2.3-fold of basal activity (Fig. 6C, S5E). Together, these data suggest that mGluR2 is activated during moderate synaptic stimulation patterns, in line with an earlier study suggesting use-dependent activation of group II mGluRs (Scanziani *et al*, 1997). Surprisingly, for all patterns of synaptic activity, levels of mGluR2 activation were almost identical next to the release site (r = 5 nm) and at the edge of the active zone (r = 100 nm) and only slowly decreased with increasing distance from the active zone (r > 100 nm, Fig. 6C, E). These results suggest that mGluR2 is efficiently activated, even at further distances from the release site, and its activation is only loosely coupled to release site location. This finding is in line with the localization of mGluR2 along the axon and inside the presynaptic bouton but not inside the active zone.

In contrast, mGluR7, having a distinctively low affinity for glutamate, was not efficiently activated by the release of a single SV, even when positioned close to the release site. At r = 5 nm, we found less than 0.3% change in activation compared to basal receptor activity (Fig. 6D, S5G). The release of 10 SVs at 5 Hz caused a relatively small increase (∼ 1.5%) in mGluR7 activity (Fig. 6D, S5H). However, the fusion of the same number of SVs at higher frequency (20 Hz) almost doubled mGluR7 response to glutamate (∼ 2.6% increase of G_αGTP_ concentration at r = 5 nm, Fig. 6D, S5I) suggesting that the level of mGluR7 activation strongly depends on the frequency of release and the peak of maximal glutamate concentration in the cleft. Additionally, the activity profiles of mGluR7 further away from the release site showed a reduction in mGluR7 response indicating that mGluR7 activation is most likely to occur in locations close to release sites (Fig. 6D, E). Altogether, these data indicate that mGluR7 is involved in the modulation of synaptic transmission only during repetitive, high-frequency release and its localization at the active zone close to the release site is curtail for its function.

A subset of parameters in the cTCAM model describing the interaction between GPCR and G-protein could be different between cell types and between the different receptors. However, there is only limited experimental data available. Therefore, we tested the influence of parameter changes on the cTCAM model output and our conclusions. We solved the model with different numerical values for the receptor activation rate, the receptor – G protein association rate, the receptor – G protein collision efficiency, and for the receptor – G protein binding affinity (Table S1). All parameter values were in the ranges previously reported (Kinzer-Ursem & Linderman, 2007). For all tested parameters, we found a similar dependency of mGluRs activation on the distance from the release site (Fig. S5F, J). However, we found that receptor – G protein collision efficiency has a higher impact on the mGluR2 and mGluR7 activation profiles.

**Figure S5.**
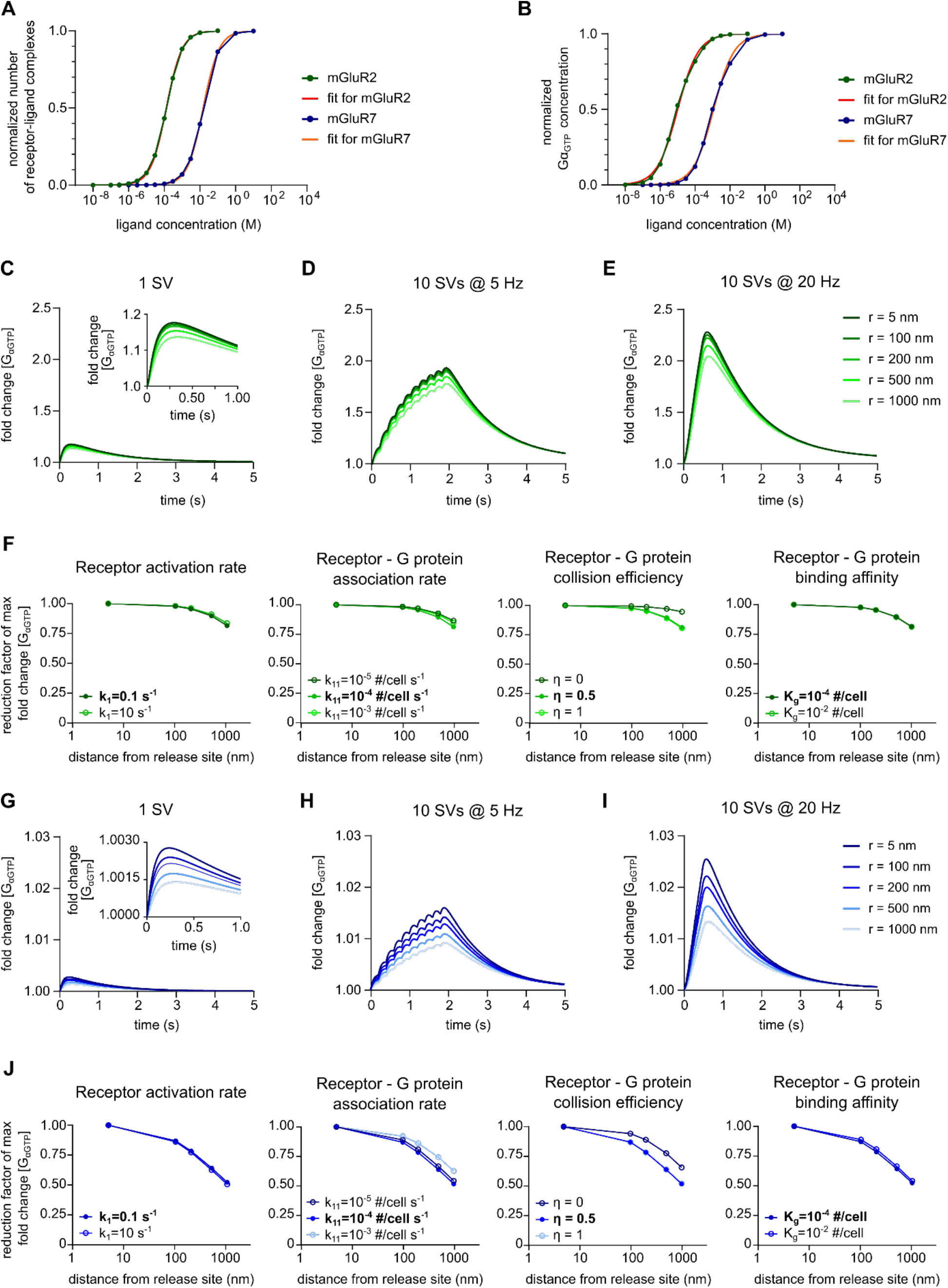
(Related to Figure 6): mGluR2 activation is loosely coupled to the distance to the release site, while mGluR7 activation is most prominent close to the release site. **(A – B)** Calibration of the model to match the number of receptor-ligand complexes **(A)** and Gα_GTP_ concentration **(B)** with reported EC_50_ values for mGluR2 and mGluR7.**(C - E)** Time courses of mGluR2 response to glutamate after the release of 1 SV **(C)**, 10 SVs at 5 Hz **(D)** and 10 SVs at 20 Hz **(E)** at different distances from the release site. **(F)** Reduction factor of mGluR2 activation at different distances from the release site after release of 10 SVs at 20Hz for various values of different parameters in cTCAM. Bold value – values used in the main model. Note that after changing one cell-specific parameters, the model was calibrated to match the reported EC_50_ values for mGluR2. **(G - I)** Time courses of mGluR7 response to glutamate after the release of 1 SV **(G)**, 10 SVs at 5 Hz **(H)** and 10 SVs at 20 Hz **(I)** at different distances from the release site. **(F)** Reduction factor of mGluR7 activation at different distances from the release site after release of 10 SVs at 20Hz for various values of different, cell-specific parameters in cTCAM. Bold value – values used in the main model. Note that after changing one cell-specific parameter, the model was calibrated to match the reported EC_50_ values for mGluR7.

## DISCUSSION

Despite the functional importance of presynaptic mGluRs in modulating the efficacy of synaptic transmission, the mechanisms that control their dynamic distribution at excitatory synapses remain poorly understood. Here, we provide new insights into the molecular mechanisms that determine the spatial distribution and mobility of presynaptic mGluRs in hippocampal neurons (Fig. 6F). We observed that presynaptic mGluR subtypes display striking differences in their subsynaptic localization and dynamics that are controlled by distinct structural mechanisms. We found that the intracellular domain of mGluR2 controls its mobility and that the actin cytoskeleton contributes to the regulation of mGluR2 diffusion behavior. In contrast, we found that the extracellular domain of mGluR7 is critical for immobilization of the receptor at presynaptic sites. Finally, a computational model of receptor activation showed that mGluR2 activation is only loosely coupled to release site location. In contrast, even when placed immediately next to the release site, there is only modest activation of mGluR7 by physiologically relevant synaptic stimulation patterns.

Mapping the precise distribution of presynaptic mGluRs is essential for understanding how these receptors contribute to synaptic transmission. In particular, the location relative to the release site is predicted to influence the probability of receptor activation and ability to trigger local downstream effectors. We found that while mGluR2 was distributed along the axon and in synaptic boutons it was largely excluded from the active zone. In contrast, we found that mGluR7 was highly enriched at the presynaptic active zone, close to the release site of synaptic vesicles. This is in line with earlier immuno-EM studies that showed that mGluR2 is present in the preterminal part of axons, but rarely found in boutons (Shigemoto *et al*, 1997), and that group III mGluRs, including mGluR7, is almost exclusively localized in the presynaptic active zone (Siddig *et al*, 2020; Shigemoto *et al*, 1997, 1996). Interestingly, these differences in localization were reflected in the surface diffusion behavior of these receptors. mGluR2 was highly mobile throughout the axon and within boutons, similar to other presynaptic receptors such as the cannabinoid type 1 receptor (CB1R) (Mikasova *et al*, 2008) and the mu-type opioid receptor (MOR) (Jullié *et al*, 2020). In contrast to these mobile receptors, however, diffusion of mGluR7 was almost exclusively restricted to presynaptic boutons. Such differences in the distribution of presynaptic receptors are likely associated with their function and may provide a means for synapses to spatially and temporally compartmentalize receptor signaling.

The differences in the distance of these mGluR2 and mGluR7 to the release site imply that these receptors respond differentially to synaptic activity. Indeed, our computational modeling studies indicate that mGluR2 activation is only loosely coupled to release site location, while mGluR7 activation is limited, even when placed in immediate proximity to the release site. These two receptor types might thus encode different modes of synaptic activity patterns: mGluR2 responding to lower frequency stimulation patterns, and mGluR7 being activated only during intense, high-frequency synaptic stimulation. It has been suggested that group III mGluRs act as auto-receptors during repetitive stimulations and modulate release probability (Billups *et al*, 2005; Pinheiro & Mulle, 2008). On the other hand, it has been described that mGluR7 is constitutively active (Kammermeier, 2015; Dunn *et al*, 2018; Stachniak *et al*, 2019), and that activity of group III mGluRs is regulated by the transsynaptic interaction with ELFN1/2 at excitatory synapses (Dunn *et al*, 2019a; Stachniak *et al*, 2019). Allosteric modulation of mGluR7 by ELFN1/2 could thus alter the threshold for receptor activation or increase its basal activity. Moreover, in our model, we assumed a homogenous distribution of G-proteins inside the presynaptic bouton and we found that changes in receptor – G protein collision efficiency had a significant impact on mGluR activation level. Therefore, we cannot exclude the possibility that at the active zone there is a higher local concentration of Gα, or that mGluR7 has a higher affinity for G-proteins than mGluR2. Thus, activation of mGluR7 could result in stronger activation of downstream signaling pathways and a larger effect on synaptic transmission. Nevertheless, the results from our computational model indicate that mGluR7 positioning relative to the release site is a critical factor increasing the probability of receptor activation.

The spatial segregation of mGluRs in presynaptic boutons could also be a mechanism to compartmentalize the downstream effectors of these receptors. Both mGluR2 and mGluR7 couple to inhibitory Gα_i_ proteins that repress adenylyl cyclase activity, decreasing cAMP production. Indeed, these receptors have overlapping downstream signaling proteins such as PKA and PKC, and are both described to modulate calcium channel activity (Martín *et al*, 2007; Ferrero *et al*, 2013; Robbe *et al*, 2002; de Jong & Verhage, 2009). But, mGluR7 has also been suggested to interact with several other components of the active zone, such as RIM1a (Pelkey *et al*, 2008), and Munc-13 (Martín *et al*, 2010). The selective effects of these receptors might thus be explained by their segregated distribution. One of the principal mechanisms of synaptic depression that is shared by these receptors, involves the interaction between the membrane-anchored βγ subunits of the G-protein with voltage-gated Ca^2+^ channels (VGCC) (Niswender & Conn, 2010; Kammermeier, 2015). An important rate-limiting factor in this mechanism is probably the distance between the G_βγ_ subunits and VGCCs. It could thus be envisioned that the effect of mGluR2 activation on synaptic transmission would not be instantaneous but would be delayed by the diffusion time of βγ subunits to VGCCs enriched at the active zone. For mGluR7 on the other hand, being immobilized in close proximity to release sites, the inhibition of VGCCs might occur much more instantaneously after receptor activation. Altogether, our data indicate that the specific modulatory effects of presynaptic mGluRs on synaptic transmission are in large part determined by their differential localization relative to the release site and their distinct surface diffusion properties.

Given the distinct distribution and diffusion properties of mGluR2 and mGluR7, we speculated that distinct mechanisms control the surface mobility of these receptors. Both C-terminal regions of mGluR2 and mGluR7 contain PDZ binding motifs, but of different types, mGluR2 contains a class I, and mGluR7 a class II binding motif (Hirbec *et al*, 2002) indicating specific intracellular interaction for each of presynaptic mGluRs. Our data indeed suggest that intracellular interactions mediated by the C-terminal region of mGluR2 regulate receptor diffusion but are not dependent on PDZ binding motifs. Instead, we found that the mobility of mGluR2 is increased by depolymerization of the actin cytoskeleton. The striking periodic organization of the actin cytoskeleton is proposed to control the distribution of receptors in the axons (Zhou *et al*, 2019) but the protocol of actin depolarization used in our single-molecule tracking experiments should not cause loss of the ring structures (Zhong *et al*, 2014). Therefore, we propose that ICD of mGluR2 introduces steric hinders for receptor diffusion. However, little is known about the molecular interaction of the C-terminal region of mGluR2 and we cannot exclude a possible direct or indirect molecular link between mGluR2 and actin. Thus, molecular mechanisms that control mGluR2 diffusion remains to be elucidated.

Also for mGluR7, it has been suggested that stable surface expression and clustering in presynaptic boutons is controlled by the intracellular interaction with the PDZ-domain, containing scaffold protein PICK1 (Boudin *et al*, 2000; Suh *et al*, 2008). In contrast, another study showed that the synaptic distribution of a mGluR7 mutant lacking the PDZ binding motif was unaltered (Zhang *et al*, 2008). Our findings suggest that the intracellular domain of mGluR7 does not have a large contribution to receptor clustering and immobilization at presynaptic boutons are consistent with this, further suggesting that interactions with PICK1 could be important for mGluR7 function but do not instruct receptor localization. Only replacing the TMD of mGluR7 with the TMD of mGluR2, but not mGluR1, caused changes in mGluR7 mobility indicating that the TMD of mGluR7 does not play a major role in controlling receptor dynamics. Based on a recent paper showing a predominant role for the TMD in mGluR2 dimerization (Gutzeit *et al*, 2019), we speculate that the increased mobility of the mGluR7 chimera containing the mGluR2 TMD could be caused by heterodimerization of this chimeric receptor with the highly mobile endogenous mGluR2. Rather, we found an unexpected role of the extracellular domain of mGluR7 in its immobilization at the presynaptic plasma membrane. Chimeric mGluR7 variants with substituted ECDs displayed higher diffusion coefficients than wild-type mGluR7 and surface diffusion was no longer restricted to the presynaptic bouton but was virtually unrestricted along the axon. Our data thus suggest that extracellular interactions can efficiently cluster the receptor and that the extracellular domain of mGluR7 is essential for immobilizing and concentrating the receptor at active zones.

The dramatic effect of replacing the extracellular domain of mGluR7 on localization and diffusion suggests that transsynaptic interactions might effectively concentrate mGluR7 at synaptic sites. This is consistent with the emerging notion that transcellular interactions greatly impact GPCR biology (Dunn *et al*, 2019a). Specifically for group III mGluRs, interactions with the adhesion molecules ELFN1 and ELFN2 have been found to modulate the functional properties of these receptors and potently impact synaptic function (Tomioka *et al*, 2014; Dunn *et al*, 2019b, 2018; Stachniak *et al*, 2019). Thus, the transsynaptic interaction with ELFN1/2 could also play a role in anchoring mGluR7 at specific synaptic sites while simultaneously regulating receptor activity via allosteric modulation. However, contribution of the transynaptic interactions of mGluR7 to its immbolization at the actize zone remains to be futher investigated.

Previous studies have suggested that ligand-induced GPCR activation, alters their surface diffusion and oligomerization properties (Yanagawa *et al*, 2018; Sungkaworn *et al*, 2017; Kasai & Kusumi, 2014; Calebiro *et al*, 2013). In heterologous cells, the diffusion rate of many GPCRs, including mGluR3 for instance, is significantly reduced after agonist stimulation (Yanagawa *et al*, 2018). Surprisingly, our data in neurons indicate that the surface mobility of mGluRs is not altered by agonist-induced receptor activation or acute increase in neuronal activity. Diffusion in the plasma membrane of heterologous cells is likely influenced by other factors than in neuronal membranes. Most notably, the unique membrane composition and expression of cell-type specific interaction partners in neurons are likely to differentially tune the diffusional properties of individual receptors. Indeed, the mobility of the CB1R in the axon decreases after desensitization (Mikasova *et al*, 2008), while the mobility of another GPCR, MOR does not change after agonist stimulation (Jullié *et al*, 2020). Our data indicate that for the presynaptic mGluRs, mGluR2, and mGluR7, structural factors, such as interactions with intra-and extracellular components predominantly instruct receptor localization, and that these mechanisms act independently of the receptor activation status. This has potentially important implications for the contribution of these receptors to the regulation of synaptic transmission. mGluR7 is likely to exert its effects very locally, restricted to individual synapses. For mGluR2 on the other hand, it could be speculated that the unchanged, high surface mobility of mGluR2 after activation allows the receptor to activate downstream effectors over larger areas, as has been suggested for the opioid receptor (Jullié *et al*, 2020). This would imply that, once activated, mGluR2 could spread its effects to neighboring synapses and dampen transmission much more globally than mGluR7 does. We can of course not exclude that only a small, undetectable subpopulation of activated mGluRs is immobilized at specific locations, but given that the threshold for mGluR2 activation is relatively low, it seems likely that the effects of mGluR2 activation are much more widespread than mGluR7. This could also imply that the activity of mGluR2 not only modulates synaptic transmission, but perhaps also controls other axonal processes such as protein synthesis, cargo trafficking, or cytoskeleton reorganization. On the other hand, after an acute decrease in neuronal activity, presynaptic mGluRs displayed lower diffusion, similar to type-A GABA receptors (Bannai *et al*, 2009). Reduction of mGluR mobility after inhibition of network activity but not after direct inhibition of receptor activity with antagonists suggests that their diffusion is not regulated by the activation status of the receptor but rather by indirect changes in membrane properties or perhaps rearrangements in scaffold proteins.

In conclusion, we identified novel regulatory mechanisms that differentially control the spatial distribution and dynamics of presynaptic glutamate receptors, that have important implications for how these receptors can contribute to the modulation of synaptic transmission. The co-existence of various other and distinct receptor types at presynaptic sites likely provides flexibility and allows synapses to differentially respond to incoming stimulation patterns. The emerging notion that different mGluRs are often co-expressed in neurons and can form functional heterodimers, further diversifies the potential of these receptors in regulating neuronal physiology. Defining the molecular mechanisms that control the dynamic spatial distribution of these receptors will be important to further our understanding of synaptic modulation.

## MATERIALS AND METHODS

### Animals

All experiments that required animals were approved by the Dutch Animal Experiments Committee (Dier Experimenten Commissie [DEC]). All animals were treated in accordance with the regulations and guidelines of Utrecht University, and conducted in agreement with Dutch law (Wet op de Dierproeven, 1996) and European regulations (Directive 2010/63/EU).

### Primary rat neuronal culture and transfection

Dissociated hippocampal cultures from embryonic day 18 (E18) Wistar rat (Janvier Labs) brains of both genders were prepared as described previously (Cunha-Ferreira *et al*, 2018). Neurons were plated on 18-mm glass coverslips coated with poly-L-lysine (37.5 mg/ml, Sigma-Aldrich) and laminin (1.25 mg/ml, Roche Diagnostics) at a density of 100,000 neurons per well in a 12-well plate. Neurons were growing in Neurobasal Medium (NB; Gibco) supplemented with 2% B27 (Gibco), 0.5 mM L-glutamine (Gibco), 15.6 µM L-glutamic acid (Sigma), and 1% penicillin/streptomycin (Gibco). Once per week, starting from DIV1, half of the medium was refreshed with BrainPhys neuronal medium (BP, STEMCELL Technologies) supplemented with 2% NeuroCult SM1 supplement (STEMCELL Technologies) and 1% penicillin/streptomycin (Gibco). Neurons were transfected at DIV3-4 (knock-in constructs) or DIV10-11 (overexpression constructs) using Lipofectamine 2000 reagent (Invitrogen). Shortly before transfection, neurons were transferred to a plate with fresh NB medium with supplements. Next, a mixture of 2 µg of DNA and 3.3 µl of Lipofectamine in 200 µl of NB medium was incubated for 15 - 30 min and added to each well. After 1 - 2 h, neurons were briefly washed with NB medium and transferred back to the plate with conditioned medium. All experiments were performed using neurons at DIV21-24.

### Antibodies and reagents

In this study the following primary antibodies were used: mouse anti-Bassoon (1:500 dilution, Enzo, #ADI-VAM-PS003-F, RRID AB_10618753); rabbit anti-GFP (1:2000 dilution, MBL Sanbio, #598, RRID AB_591819); rabbit anti-mGluR2/3 (1:50 dilution, EMD Millipore, #AB1553, RRID AB_90767); rabbit anti-mGluR7 (1:100 dilution, Merck Millipore, #07-239, RRID AB_310459); and anti-GFP nanobodies conjugated with ATTO647N (1:15000 dilution, GFPBooster-ATTO647N, Chromotek, #gba647n). The following secondary antibodies were used: goat Abberior STAR580-conjugated anti-rabbit (1:200 dilution, Abberior GmbH, #2-0012-005-8) and goat Abberior STAR635P-conjugated anti-mouse (1:200 dilution, Abberior GmBH, #2-0002-007-5). The following chemical reagents were used: 4-aminopyridine (4-AP, TOCRIS, #940), ADX 71743 (TOCRIS, # 5715), DL-TBOA (TOCRIS, #1223), L-AP4 (TOCRIS, #0103), Latrunculin B (Bio-Connect, # SC-203318), LY379268 (TOCRIS, #2453), (RS)-APICA (TOCRIS, # 1073) and Tetrodotoxin citrate (TTX, TOCRIS, #1069).

### DNA plasmids

The SEP-mGluR2 CRISPR/Cas9 knock-in constructs were designed as described in (Willems *et al*, 2020). SEP tag was inserted into exon 2 of *Grm2* gene using following target sequence: 5’-AGGGTCAGCACCTTCTTGGC-3’. Plasmids pRK5-mGluR2-GFP and pRK5-myc-mGluR7a (gift from Dr. J. Perroy) were used as PCR template to generate pRK5-SEP-mGluR2 and pRK5-SEP-mGluR7. pRK5-mOrange-mGluR2 and pRK5-mOrange-mGluR7 were created by exchanging SEP with mOrange in pRK5-SEP-mGluR2 and pRK5-SEP-mGluR7. pRK5-SEP-mGluR7-N74K was cloned using site-directed mutagenesis with the following primers: forward: 5’-GGCGACATCAAGAGGGAGAAAGGGATCCACAGGCTGGAAGC-3’ and reverse: 5’-GCTTCCAGCCTGTGGATCCCTTTCTCCCTCTTGATGTCGCC-3’. To create SEP-tagged chimeric variants of mGluR2 and mGluR7, sequences of wild-type receptors in pRK5-SEP-mGluR2 and pRK5-SEP-mGluR7 were replaced by the sequence of the chimeric receptor. Chimeric receptors were cloned by fusing sequences encoding different domains of mGluR2, mGluR7 and mGluR1 as follow:

mGluR2-ICD7: 1-819 aa mGluR2 + 849-913 aa mGluR7;

mGluR2-TMD7: 1-556 aa mGluR2 + 578-848 aa mGluR7 + 820-872 mGluR2;

mGluR2-ECD7: 1-583 aa mGluR7 + 562-872 aa mGluR2;

mGluR7-ICD2: 1-848 aa mGluR7 + 820-872 aa mGluR2;

mGluR7-TMD1: 1-588 aa mGluR7 + 591-839 aa mGluR1+ 849-914 aa mGluR7;

mGluR7-TMD2: 1-588 aa mGluR7 + 568-819 aa mGluR2 + 849-914 aa mGluR7;

mGluR7-ECD1: 1-585 aa mGluR1 + 584-913 aa mGluR7;

mGluR7-ECD2: 1-556 aa mGluR2 + 584-913 aa mGluR7.

Amino acid numbering is based on sequences in UniPortKB database (mGluR1 - Q13255-1, mGluR2 - P31421-1, mGluR7 - P35400-1) and starts with the first amino acid of the signal peptide. pRK5-SEP-mGluR1 (Scheefhals *et al*, 2019) was used as a PCR template for the transmembrane and extracellular domain of mGluR1. pRK5-SEP-mGluR2-ΔICD (lacing ICD, 823 - 872 aa) and pRK5-SEP-mGluR2-ΔPDZ (lacing PDZ binding motif, 869 – 872 aa) were cloned using primers containing the desired mutation. All chimeric and deletion mGluR variants were cloned using Gibson assembly (NEBuilder HiFi DNA assembly cloning kit). pRK5-mOrange-mGluR2-ECD7 was generated by replacing SEP tag in pRK5-SEP-mGluR2-ECD7. Synaptophysin1-mCherry plasmid was generated by replacing the pHluorin-tag in Synaptophysin1-pHluorin (gift from L. Lagnado, Addgene plasmid # 24478, (Granseth *et al*, 2006)) with mCherry from pmCherry-N1 (Invitrogen). All sequences were verified by DNA sequencing.

### Immunostaining and gSTED imaging

Neurons at DIV21 were fixed with 4% PFA and 4% sucrose in PBS for 10 min at RT and washed three times with PBS supplemented with 100 mM glycine. Next, cells were permeabilized and blocked with 0.1% Triton-X (Sigma), 10% normal goat serum (Abcam), and 100 mM glycine in PBS for 1 h at 37°C. Neurons were incubated with primary antibodies diluted in PBS supplemented with 0.1% Triton-X, 5% normal goat serum, and 100 mM glycine for 3 - 4h at RT. After three times washing cells with PBS with 100 mM glycine, neurons were incubated with secondary antibodies diluted in PBS supplements with 0.1% Triton-X, 5% normal goat serum, and 100 mM glycine for 1h at RT. Cells were washed two times with PBS with 100 mM glycine and two times with PBS. Neurons were mounted in Mowiol mounting medium (Sigma). Due to low endogenous level of mGluR2, the signal of endogenously tagged protein was enhanced by immunostaining with rabbit anti-GFP antibodies (1:2000 dilution, MBL Sanbio). Dual-color gated STED imaging was performed with a Leica TCS SP8 STED 3 microscope using an HC PL APO 100/1.4 oil-immersion STED WHITE objective. Abberior STAR 580 and 635P were excited with 561 nm and 633 nm pulsed laser light (white light laser, 80 MHz) respectively. Both Abberior STAR 580 and 635P were depleted with a 775 nm pulsed depletion laser. Fluorescence emission was detected using a Leica HyD hybrid detector with a gating time from 0.5 ns to 6 ns.

### Live-cell imaging and FRAP experiments

For all live-cell imaging experiments, cells were kept in a modified Tyrode’s solution (pH 7.4) containing 25 mM HEPES, 119 mM NaCl, 2.4 mM KCl, 2 mM CaCl_2_, 2 mM MgCl_2_, 30 mM glucose. FRAP experiments were carried out in an environmental chamber at 37°C (TokaHit) on an inverted Nikon Ti Eclipse microscope equipped with a confocal spinning disk unit (Yokogawa), an ILas FRAP unit (Roper Scientific France/ PICT-IBiSA, Institut Curie), and a 491-nm laser (Cobolt Calypso). Fluorescence emission was detected using a 100x oil-immersion objective (Nikon Apo, NA 1.4) together with an EM-CCD camera (Photometirc Evolve 512) controlled by MetaMorph 7.7 software (Molecular Devices). Images were acquired at 1 Hz with an exposure time between 100 and 200 ms. 3 - 5 ROIs covering single boutons were bleached per field of view.

### Single-molecule tracking with uPAINT

Single-molecule tracking was carried out in modified Tyrode’s solution supplement with 0.8% BSA and ATTO647N-conjugated anti-GFP nanobodies (1:15000 dilution, GFPBooster-ATTO647N, Chromotek, #gba647n) on Nanoimager microscope (Oxford Nanoimaging; ONI) equipped with a 100x oil-immersion objective (Olympus Plans Apo, NA 1.4), an XYZ closed-loop piezo stage, 471-nm, 561-nm and 640-nm lasers used for excitation of SEP, mCherry and ATTO647N respectively. Fluorescence emission was detected using an sCMOS camera (ORCA Flash 4, Hamamatsu). 3,000 images were acquired in stream mode at 50 Hz in TIRF. Before every tracking acquisition, 30 frames of SEP and mCherry signal were taken to visualize cell morphology or boutons. To determined how the activity of receptors influences their diffusion, first control acquisitions (2 - 3 fields of view per coverslip) were taken, next chemical reagents (with final concentrations: 200 µM 4-aminopyridine (4-AP) + 10 µM DL-TBOA; 30 µM ADX; 500 µM L-AP4; 100 µM APICA; 100 µM LY379268; 5 µM Latrunculin-B; 1 µM TTX) or high K^+^ solution (2x) were added to imaging chamber, incubated for 3 - 5 min and final acquisitions of previously imaged fields of views were performed. A high K^+^ solution was prepared by replacing 45 mM NaCl with KCl. Total incubation times with chemical reagents or high K^+^ solution did not exceed 15 min.

### Computational modeling of mGluR activity

#### Receptor model

To study the time-dependent response of mGluRs upon glutamate release, a G-protein-coupled receptor model was combined with the time-dependent concentration profile of glutamate released from synaptic vesicles. The cubic ternary complex activation model (cTCAM) of GPCR signaling describes the interaction of the receptors *R*, ligands *L* and G-proteins *G* (Kinzer-Ursem & Linderman, 2007). The receptors can complex with G-proteins to form *RG* and furthermore, can be in an active state *R*^*^ denoted by the asterisk. G proteins are produced by a cascade of *Gα*_*GTP*_ hydrolysis and *G*_*βγ*_ binding. The reactions are described by the following differential equations:

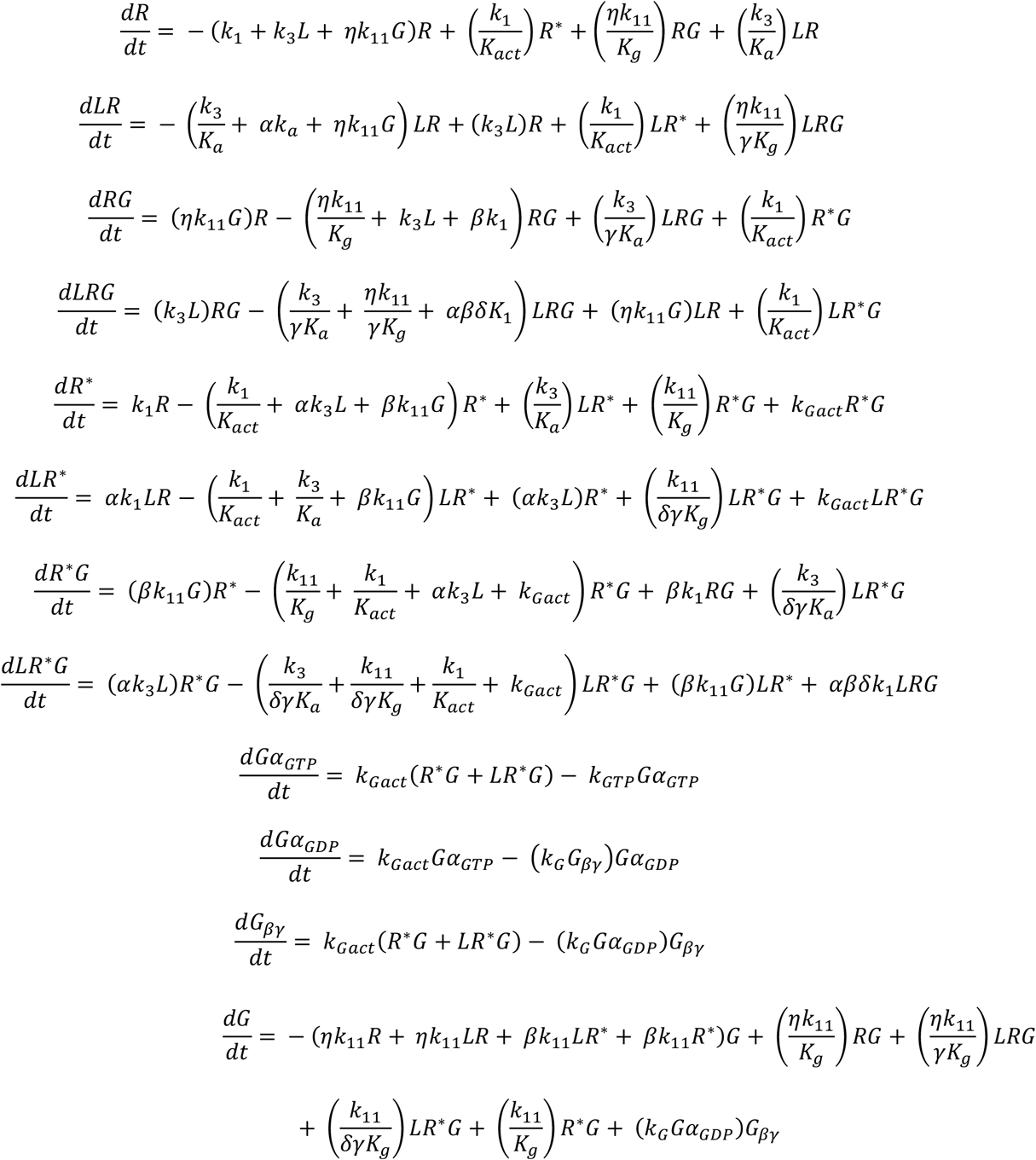

To find the steady-state solution without ligand (*L* = 0), these equations were solved with initial conditions *R* = 100, *G* = 1000, and the remaining variables set to zero using the NDSolve function of Mathematica (version 12.0, Wolfram Research Inc.). The numerical values for the used parameters have been described previously (Kinzer-Ursem & Linderman, 2007) and are summarized in Table S1. The number of receptors and G-proteins in presynaptic bouton are estimated based on quantitative mass-spectrometry data published in (Wilhelm *et al*, 2014). To describe the different behaviors of mGluR2 and mGluR7, only the association constant *K*_*a*_ was adjusted to match previously published EC_50_ values: 10 µM for mGluR2 and 1 mM for mGluR7 (Schoepp *et al*, 1999). The EC_50_ value is the concentration of the ligand that gives the half-maximum response. Hence, the response was estimated by the number of *Gα*_*GTP*_. The steady-state solution without ligand was used as the initial state of the system and the new steady-state values for different amounts of the ligand were numerically determined. The relative normalized change of *Gα*_*GTP*_ gives the response:

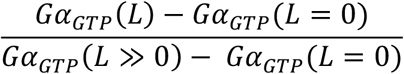

To obtain the EC_50_ value, the following function was fitted to the data points from the numerical solution (Figure S6B):

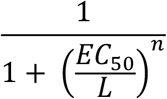

In this way, a parameterization of mGluR2 with *K*_*a*_ = 0.7·10^4^ M^-1^ and respective EC_50_ = 10 µM, and mGluR7 with *K*_*a*_ = 60 M^-1^ and respective EC_50_ = 1.15 mM was obtained. To investigate the ligand-receptor affinity, the normalized response of the sum of all formed receptor-ligand complexes was determined as (Figure S6A):

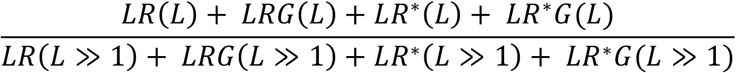

#### Diffusion model

The time-dependent concentration of glutamate released from a synaptic vesicle was described as a point source on an infinite plane. The solution of the diffusion equation gives the surface density:

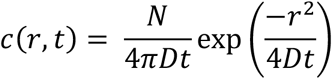

in which *r*: the distance from the source,

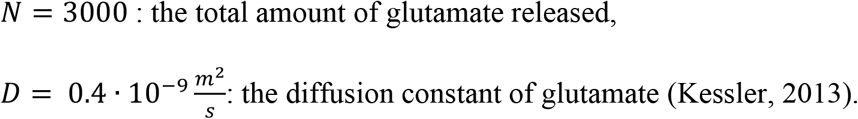

To transform the surface density into a concentration the following formula was used:

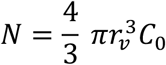

in which *r*_*v*_ = 25 *nm*: the radius of a vesicle,

*C*_0_: the glutamate concentration inside the vesicle.

Next, the surface density was divided by the *d* = 20 *nm* width of the synaptic cleft to obtain:

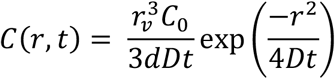

Hence, the initial concentration is given by:

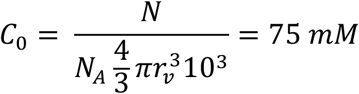

in which *N*_*A*_: Avogadro’s constant.

To describe the glutamate concentration from a sequence of vesicles release events, superposition was used as follows:

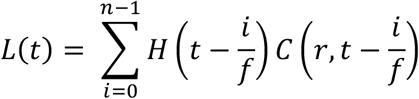

in which *n*: the number of vesicles released,

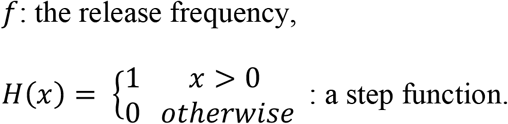

The diffusion profile was combined with the receptor model and the differential equations were solved numerically for a given distance *r* from the release site. For the initial conditions, the steady-state solution without ligand was used. Because of the non-linearities in the equations and the possible large values of the concentration profile for small times, to solve the equations numerically, we reduced the accuracy and precision of the numerical integration method in Mathematica’s NDSolve function. This adjustment potentially introduced an error of less than 5%, which is small enough to be neglected in our analysis and conclusions.

#### Testing the model with different sets of parameter

To test the model’s output sensitivity to the numerical values of the model parameters, parameters: the receptor activation rate, the G-protein association rate, the G-protein collision efficiency, and the G-protein binding affinity were changed in the ranges reported previously (Kinzer-Ursem & Linderman, 2007). After changing the value of one of the parameters, the K_a_ was always adjusted to match the EC_50_ values of the specific receptors. Note, the nonlinearity and the multiple pathways of the model cause also changes in K_a_ values in some cases (Table S1). Next, the diffusion profile of glutamate released from 10 SV at 20Hz was combined with the receptor models described by different sets of parameters. A change of the numerical values for the parameter changed the absolute value of the model output, but the overall trend of receptor activation remained the same. For better comparison of how changes in parameter values influence receptor activation at different distances from the release site, the maximum fold changes of G_αGTP_ concentration for each receptor at different distances were normalized to the maximum of this concentration at 5 nm away from the release site to obtain the reduction factor of the maximum fold change of G_αGTP_ concentration (Figure S6F, J).

**Table S1.**
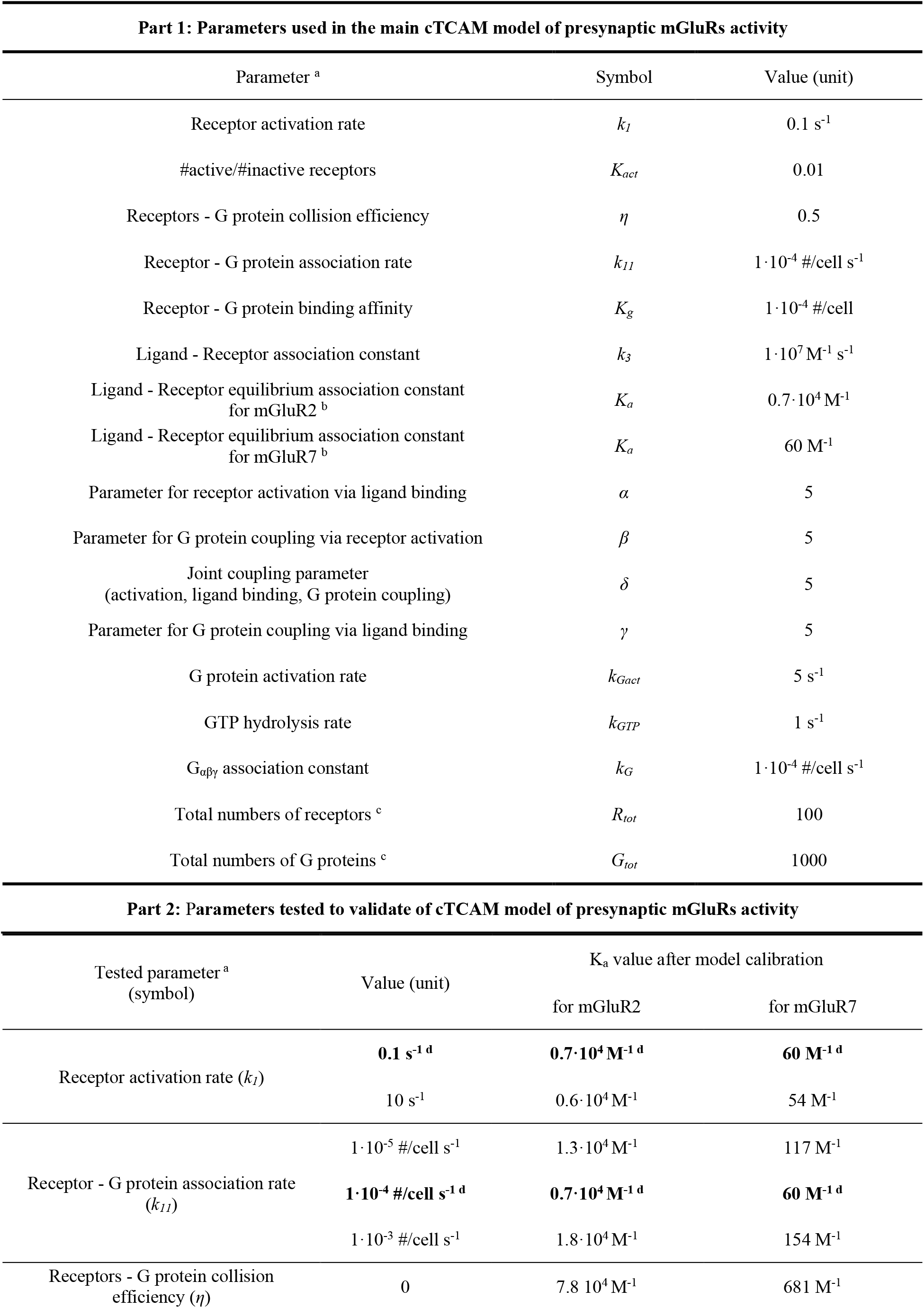
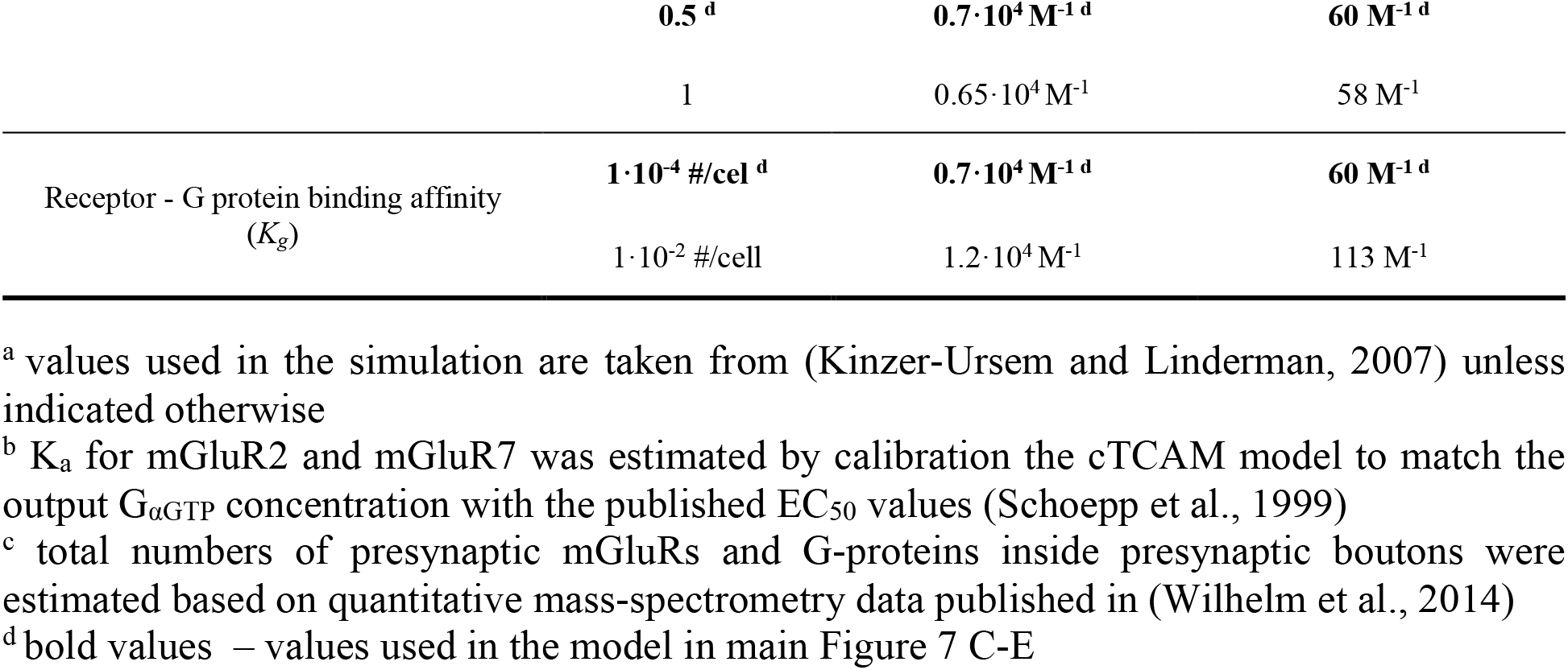
(Related to Figure 6): Parameters used in the cTCAM model of presynaptic mGluRs activity.

### Quantification of co-localization

Analysis of co-localization between Bsn and mGluRs was done using Spot Detector and Colocalization Studio plug-ins built-in in Icy software (De Chaumont *et al*, 2012). Objects detected with Spot Detector (size of detected spots: ∼7 pixel with sensitivity 100 and ∼13 pixels with sensitivity 80) were loaded into Colocalization Studio and statistical object distance analysis (SODA) (Lagache *et al*, 2018) was performed to obtain the fraction of mGluR spots co-localized with Bsn spots.

### Quantification of SEP-mGluRs bouton enrichment

Neuron co-expressing cytosolic mCherry and SEP-mGluR2 or SEP-mGluR7 were fixed at DIV21 with 4% PFA and 4% sucrose from 10 min in RT. Next, cells were washed three times with PBS and mounted in Mowiol mounting medium (Sigma). Imaging was performed with Zeiss LSM 700 confocal microscope using 63× NA 1.40 oil objective. To analyze the enrichment of mGluRs in presynaptic boutons, line profiles along boutons and neighboring axonal region were drawn in ImageJ (line width 3 pixels). Next, intensity profiles were fitted with a Gaussian function in GraphPad Prism. To calculate the ratio of intensity in bouton over axon, the amplitude of the Gaussian fit was divided by the minimum value of the fit.

### Quantification of the expression level of SEP-mGluRs

Neurons transfected with SEP-mGluR2 or SEP-mGluR7 at DIV 10 were fixed and immuno-stained at DIV21. To compare expression levels of SEP-tagged mGluRs with the endogenous levels, neurons expressing SEP-mGluR2 were co-stained with anti-mGluR2/3, and cells expressing SEP-mGluR7 were co-stained with anit-mGluR7. Imaging was performing on a Leica TCS SP8 STED 3 microscope using an HC PL APO 100/1.4 oil-immersion STED WHITE objective keeping all imaging parameters constant. In ImageJ, circular ROIs with the size of boutons were drawn around boutons in neurons expressing SEP-tagged mGluRs and untransfected boutons positive for mGluR2 or mGluR7. Fluorescent intensity values of control boutons (untransfected) and expressing SEP-tagged mGluRs were normalized by dividing by the average intensity values of all control boutons.

### Quantification of FRAP experiments

Time series obtained during FRAP experiments were corrected for drift when needed using Template Matching plug-in in ImageJ. Circular ROIs with the size of the bleached area were drawn in ImageJ. Fluorescent intensity transients were normalized by subtracting the intensity values of the 1^st^ frame after bleaching and dividing by the average intensity value of the baseline (5 frames before bleaching). The mobile fraction was calculated by averaging the values of the last 5 points of fluorescent transients. τ of recovery was determined by fitting a single exponential function to the recovery traces.

### Single-molecule tracking analysis

NimOS software (Oxford Nanoimager; ONI) was used to detect the localization of single molecules in uPAINT experiments. Molecules with a localization precision < 50 nm and photon count > 200 photons were used for analysis. To filter out unspecific background localizations from outside neurons, a cell mask based on the SEP image was created using an adaptive image threshold in Matlab (sensitivity 40-55). Only localizations inside the mask were selected for further analysis. Tracking and calculation of the diffusion coefficient were performed in custom-written Matlab (MathWorks) scripts described previously (Willems *et al*, 2020). Only trajectories longer than 30 frames were used to estimate the instantaneous diffusion coefficient. Classification of molecule state as mobile or immobile was based on the ratio between the radius of gyration and mean step size of individual trajectories using formula 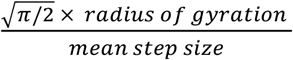 (Golan & Sherman, 2017) (Figure S2F-H). Molecules with a ratio < 2.11 were considered immobile (Golan & Sherman, 2017). A mask of presynaptic boutons was created based on the TIRF image of Synaptophysin1-mCherry as previously described (Li & Blanpied, 2016). Synaptic trajectories were defined as trajectories that had at least one localization inside the bouton mask. In all graphs representing D_eff_ or mobile fraction, a single data point is an average D_eff_ or mobile fraction from one complete field of view.

### Statistical analysis

All used in this study statistical tests are described in figure legends and the main text. All statistical analyses and graphs were prepared in GraphPad Prism. Figures were created in Inkscape.

## AUTHOR CONTRIBUTIONS

Conceptualization, Methodology, Validation, & Formal Analysis, A.B., F.B. and

H.D.M.; Investigation, A.B. and F.B.; Resources,

H.D.M.; Writing – Original Draft & Editing, A.B. and

H.D.M.; Writing – Review, F.B.; Visualization, A.B.;

Supervision, H.D.M.; Funding Acquisition, H.D.M.

## ACKNOWLEDGEMENTS

We would like to thank dr. Arthur de Jong for critical reading of the manuscript, Manon Westra for help with Matlab scripts, and all members of the MacGillavry lab for helpful discussions. This work was supported by the European Research Council (ERC-StG 716011) to H.D.M.

## DECLARATION OF INTERESTS

The authors declare no competing interests.

## Notes

### Competing Interest Statement

The authors have declared no competing interest.

